# TRIP13 protects pancreatic cancer cells against intrinsic and therapy-induced DNA replication stress

**DOI:** 10.1101/2025.01.26.634889

**Authors:** Jay R. Anand, Gaith N. Droby, Sayali Joseph, Urvi Patel, Xingyuan Zhang, Jeffrey A. Klomp, Channing J. Der, Jeremy E. Purvis, Samuel C. Wolff, Jessica Bowser, Cyrus Vaziri

## Abstract

Oncogene activation in normal untransformed cells induces DNA replication stress and creates a dependency on DNA Damage Response (DDR) mechanisms for cell survival. Different oncogenic stimuli signal via distinct mechanisms in every cancer setting. The DDR is also pathologically re-programmed and deployed in diverse ways in different cancers. Because mutant KRAS is the driver oncogene in 90% of Pancreatic Ductal Adenocarcinomas (PDAC), here we have investigated DDR mechanisms by which KRAS-induced DNA replication stress is tolerated in normal human pancreatic epithelial cells (HPNE). Using a candidate screening approach, we identify TRIP13 as a KRAS^G12V^-induced mRNA that is also expressed at high levels in PDAC relative to normal tissues. Using genetic and pharmacological tools, we show that TRIP13 is necessary to sustain ongoing DNA synthesis and viability specifically in KRAS^G12V^-expressing cells. TRIP13 promotes survival of KRAS^G12V^-expressing HPNE cells in a Homologous Recombination (HR)-dependent manner.

KRAS^G12V^-expressing HPNE cells lacking TRIP13 acquire hallmark HR-deficiency (HRD) phenotypes including sensitivity to inhibitors of Trans-Lesion Synthesis (TLS) and Poly-ADP Ribose Polymerase (PARP).

Established PDAC cell lines are also sensitized to intrinsic DNA damage and therapy-induced genotoxicity following TRIP13-depletion. Taken together our results expose TRIP13 as an attractive new and therapeutically-tractable vulnerability of KRAS-mutant PDAC.

## INTRODUCTION

For decades it has been recognized that strong mitogenic stimuli can be cytostatic. For example excessive growth factor signaling (1–6) or overproduced oncoproteins (including c-MYC (7–13), Cyclin E (11,14–17), β- catenin (18), and active mutants of KRAS (19–21)) arrest the cell cycle. The cytostatic effects of excessive mitogenic stimulation are often attributed to DNA replication stress and DNA damage. Many mechanisms are proposed for why oncogenic stimuli induce replication stress including: re-firing of DNA replication origins more than once every S-phase, accelerated G1 entry leading to reduced DNA replication licensing, Reactive Oxygen Species (ROS), cohesion complex occupancy, R-loops and Transcription Replication Collisions (TRC) (22–25). The DNA Damage Response (DDR) mechanisms deployed to sense and remediate oncogene-associated DNA replication stress are highly consequential because they dictate cell plasticity and fate, survival, and genome stability.

We and others have previously described some of the DDR mechanisms that respond to oncogenic stimuli (14,26,27). However, past studies have often deployed artificial experimental systems to overexpress oncogenes at levels that are not pathologically-relevant. For example, massive overexpression of Cyclin E (at levels up to >10-fold higher than are expressed in cancer cells) is a standard experimental model system (14,28,29). The robust DNA replication stress response elicited by overexpressed Cyclin E is highly atypical of most oncogenic stimuli. Past studies have routinely used Cyclin E to hyper-stimulate S-phase in cancer cell lines such as U2OS which already experience intrinsic DNA replication stress and have atypical DDR properties when compared with normal cells (11,14,15,17,26,28,30). Therefore, we have a limited mechanistic understanding of how normal cells respond to pathologically relevant oncogene-associated stresses.

No single mechanism explains responses to DNA replication stresses induced by all oncogenic stimuli. Different oncogenic stimuli act via distinct mechanisms within a given cell type, and the same oncogenic stimulus can frequently elicit different responses in different cell types (31–34). Similarly, the DDR is reprogrammed and deployed in diverse ways in different cancers (35). Therefore, it is necessary to determine the relationship between specific oncogenes and the DDR in defined disease-relevant settings. There is no consensus on how oncogenic KRAS-induces DNA replication stress or how that stress is remediated. For example, it is reported that KRAS both does (36) and does not (37) induce DNA replication stress. Effects of KRAS signaling are notoriously tissue and context-specific (32,38–42) and cellular models for studying responses to KRAS must be chosen carefully to correspond to the disease model.

Oncogenic KRAS drives 90% of Pancreatic Ductal Adenocarcinomas (PDACs) (43,44). Of all major cancers, Pancreatic Ductal Adenocarcinoma (PDAC) has the highest mortality rate (45) and the 5-year survival rate of PDAC patients is a dismal 13% (46–48). Recently developed direct inhibitors of one KRAS mutant, KRAS^G12C^ (G12Ci) have shown promising responses in KRAS^G12C^ mutant PDAC, supporting the clinical potential of anti-KRAS therapies in PDAC treatment (49,50). Although KRAS inhibitors are initially effective for treating PDAC, patients usually relapse as their tumors adapt to KRAS blockade (51–54) and preclinical data support the emergence of similar resistance mechanisms in the context of PDAC (55,56). There is an urgent need to identify additional dependencies of KRAS-driven PDAC and exploit those vulnerabilities as therapeutic targets. DNA-damaging agents (e.g. gemcitabine) are standard-of-care treatments for PDAC. However, innate and acquired chemoresistance limit efficacy of genotoxic therapies (57–59). It is also critical to elucidate and target chemoresistance mechanisms if we are to curtail the high mortality from PDAC.

Previous studies have highlighted the presence of DDR markers in PDAC samples, supporting a role for oncogene-induced DNA replication stress in the etiology of pancreatic cancer (60). However, there is no paradigm for how KRAS induces DNA damage during pancreatic carcinogenesis, or how the DDR protects PDAC and their normal precursors from KRAS genotoxicity. To address this knowledge gap, we opted to study consequences of KRAS-induced mitogenesis in normal untransformed Human Pancreatic Nestin-Expressing (HPNE) cells (61). We reasoned that KRAS signaling critically regulates mitogenesis in the pancreatic epithelium (62). Furthermore, activation of KRAS by mutation in normal pancreatic epithelial cells leads to premalignant lesions termed pancreatic intraepithelial neoplasias (PanINs) (45,63). Therefore, the relationships between KRAS and DDR that we define in HPNE will likely be both physiologically and pathologically relevant.

We have developed an experimental system that allows for tight regulation of oncogenic KRAS expression while avoiding artificially-high hyper-physiological mitogenic signaling. Using this model system, we identify TRIP13 as an exquisitely-specific requirement for DNA synthesis and viability of KRAS-expressing cells.

Critically, TRIP13 is pharmacologically-tractable and provides opportunities for new therapies that target a unique dependency of KRAS-driven PDAC.

## MATERIALS AND METHODS

### Cell culture and chemical agents

hTERT-HPNE (ATCC CRL-4023) cells were purchased from the American Type Culture Collection. Pa02c, Pa03c and hTERT-immortalized HPNE-DT expressing wild type KRAS or KRAS^G12V^ were provided by Dr. Channing Der (UNC). AsPc-1, Panc10.05 and Capan-1 cell lines were purchased from the UNC tissue culture facility. Cells were grown at 37 °C in DMEM medium supplemented with 10% tetracycline negative (Tet -ve) fetal bovine serum and penicillin–streptomycin (1%), in humidified chambers with 5% CO2. Capan-1 cells were grown in IMDM medium supplemented with 20% fetal bovine serum and penicillin–streptomycin (1%). The following drugs were used in this study: doxycycline hydrochloride (Fisher scientific, BP26531), DCZ0415 (MedChemExpress, HY-130603), Olaparib (MedChemExpress, HY-10162), Gemcitabine (MedChemExpress, HY-17026), AZD1775 (MedChemExpress, HY-10993).

### LentiCRISPR and sgRNA Cloning

To generate knockout cell lines, a blasticidin S deaminase (BSD) selection marker and a destabilization domain (DD) were cloned into lentiCRISPR v2 (Addgene #52961) via NEBuilder HiFi DNA Assembly in a stepwise manner. BSD cloning was performed by assembling 3 pieces: 2 PCR-amplified pieces of lentiCRISPR v2 and 1 PCR-amplified piece of pLX304 (Addgene #25890) with 13 bp overlaps. lentiCRISPR v2 primers were used to remove puromycin N-acetyltransferase selection marker. DD cloning was performed by assembling 2 pieces: 1 PCR-amplified pieces of lentiCRISPR v2 BSD and 1 PCR-amplified piece of DD-Cas9 with filler sequence and Venus (Addgene #90085) with 25 bp overlaps. Small PCR products were purified using a PCR purification kit (QIAGEN, 28104) and large PCR products (>6kb) were run on a 1% agarose gel and fragments were extracted using a gel extraction kit (QIAGEN, 28706). Fragments were assembled by HiFi DNA Assembly according to manufacturer instructions (NEB, E2621). Electroporation: 0.75 µL of HiFi assembly mixture was added to 25 µL of electrocompetent bacteria (Lucigen, 60242-2). The bacteria and DNA mixture was electroporated in ice-chilled cuvettes (Bio-Rad, 1652083) using a Gene Pulser Xcell electroporator (Bio-Rad, 1652660) at 1800 Volts, 10 µFarad, 600 Ohm, 1 mm cuvette gap. 500 µL of recovery media was added immediately post electroporation (Lucigen, 80026-1). Transformed bacteria were incubated at 37 °C for 1 hour then plated on LB ampicillin plates and incubated overnight at 37 °C. Individual bacterial colonies were transferred to 500 mL LB cultures and incubated at 37 °C for 16 hours. Cloned plasmid DNA was extracted using plasmid maxiprep kit (QIAGEN, 12362). Oligonucleotides encoding sgRNA targeting TRIP13 gene and a non-targeting control sgRNA (see Supplementary Table 1 “Oligo sequences”) were cloned into LentiCRISPRv2 BSD DD-cas9 vector. The resulting vectors were transformed into Endura Chemically Competent Cells according to the manufacturer’s protocol.

### Lentivirus generation

To generate lentiviruses for gene knockout or knock-in, replication-incompetent lentivirus was packaged via transfection of HEK293 T cells with a either a lentiCRISPRv2 SpCas9 or LV-mCherry-FLuc (a gift from Shawn Hingtgen, UNC) vector and viral packaging plasmids, pMD2.G and psPAX2, gifts from Didier Trono (Addgene # 12259 and 12260) using a 4:1:3 ratio (12 µg DNA per transfection reaction) with Lipofectamine 2000. One day after the transfection, culture medium was changed. Lentivirus-containing culture medium from transfected cells was collected at 24 h and 48 h post medium change, filtered through 0.45 µm filter, aliquoted, and stored at −80 °C.

### Lentiviral transduction to generation of stable cell lines

To generate the doxycycline (Dox)-inducible hTERT-HPNE cell lines stably expressing HA-KRAS^G12V^, the cDNA fragment encoding HA-KRAS^G12V^ was PCR amplified from pCGN-HA-K-4B(G12V) (64) and gateway-cloned into the pinducer20 plasmid, which placed it under transcriptional control of a doxycycline-regulated promoter. High-titer lentivirus was produced in HEK293T cells and hTERT-HPNE cells were infected with lentivirus-containing medium containing 8 mg/ml polybrene (Sigma-Aldrich TR-1003-G). Medium was changed after 24 hours and stably transduced cells were selected by growth in medium containing 1000 mg /ml G418 (ThermoFisher). To avoid clonal selection of idiosyncratic cells, pools of stably-infected cells were used for all experiments. For all HPNE experiments, doxycycline was replenished every 3 days. To generate *TRIP13* knockout PDAC cell lines, cultures were transduced with lentiviruses (lentiCRISPRv2 SpCas9 sgTRIP13) then BSD-selected after overnight infection. To generate luciferase-expressing HPNE cell lines, hTERT-HPNE EV and KRAS^G12V^ were transduced with mCherry-FLuc lentiviruses then sorted on the FACSAria II cell sorter.

### 3D spheroids

2000 cells were grown in ultra-low attachment (ULA) U-botom 96-well plates(Corning, 650970) with DMEM supplemented with 10% FBS and 1% penicillin/streptomycin for 4 days. For imaging, spheroids were incubated with Hoechst 33342 (Invitrogen, R37605; 1:200; 20 mins) in 96 well plate. Propidium iodide was subsequently added to the wells at a final concentration of 8 ug/ml just before imaging. Images as a z-stack were taken using the 10X lens of the Keyence BZ-X800 Microscope and were processed using Keyence imaging software. For competitive growth assay, sgNon-Targteing and sgTRIP13 cells were mixed with H2B-GFP expressing cells in 1:1 ratio and seeded in 96-well ULA plates for a week. Every two days, half of the media was replaced with fresh media. After a week, spheroids were collected in a 15 ml tube, washed with PBS, dissociated with Accutase, strained through 40 μm nylon cell strainer (Falcon, 352340), and analyzed by flow cytometer to determine number of GFP-positive and GFP-negative cells. Ratios of GFP_positive and GFP_negative cells were quantified to analyze relative growth. For viability analysis, cells were grown in 96-well ULA plates. Next day, drug was added by replacing half of the media with fresh media containing drug. After three days of drug treatment, cell viability was assessed using the CellTiter-Glo® 3D Cell Viability Assay.

### Immunoblotting

Immunoblotting methods were carried out essentially as described (65–67). In brief, cells grown in plates were washed thrice in ice-cold PBS and lysed in 100–200 µL of ice-cold cytoskeleton buffer (CSK buffer; 10 mM Pipes, pH 6.8, 100 mM NaCl, 300 mM sucrose, 3 mM MgCl2, 1 mM EGTA, 1 mM dithiothreitol, 0.1 mM ATP, 10 mM NaF, and 0.1% Triton X-100) freshly supplemented with complete protease inhibitor cocktail (Roche, Indianapolis, IN, USA) and PhosSTOP (Roche). In the case of spheroids, they were collected in a 15 ml tube, washed with PBS and dissociated with Accutase before adding the CSK buffer. Cell lysates were centrifuged at 1500g for 4 min to remove the CSK-insoluble chromatin fraction. The detergent-insoluble chromatin fractions were washed once with 1 ml of CSK buffer, resuspended in CSK, and treated with nuclease. Cell extracts were separated by SDS-PAGE, transferred to nitrocellulose membranes, and incubated overnight with the primary antibodies and 1 hour with the secondary antibodies in 5% nonfat milk TBST. The sources of antibodies and dilutions at which they were used are as follows: KRAS (Santa Cruz Biotechnology, sc-30; 1:500); phosphor- p44/42 MAPK (Erk1/2) (Thr202/Tyr204) (Cell Signaling Technology, 9106; 1:1000); HA-Tag (Santa Cruz Biotechnology, sc-7392; 1:1000); GAPDH (Santa Cruz Biotechnology, sc-32233; 1:5000); VINCULIN (Sigma Aldrich, V4505; 1:10000); phospho-Histone H2A.X (Ser139) (Millipore Sigma, 05-636; 1:5000); PCNA (Santa Cruz Biotechnology, sc-56; 1:500); phosphor RPA32 (S33) (Bethyl, A300-246A; 1:1000); β-actin (Santa Cruz Biotechnology, sc-130656; 1:5000); CDT1 (Bethyl, A300-786A; 1:1000); MCM2 (Santa Cruz Biotechnology, sc- 10771; 1:1000); CDC45 (Santa Cruz Biotechnology, sc-9298; 1:1000); TRIP13 (Santa Cruz Biotechnology, sc- 514285; 1:500); phospho-ATM(Ser1981) (Santa Cruz Technology, sc-47739; 1:500); ZRANB3 (Bethyl, A303-033A-M; 1:1000); PRIMPOL (Proteintech, 29824-1-AP; 1:1000). Perkin Elmer Western Lightning Plus ECL was used to develop films.

### Population doubling

HPNE cells were counted, seeded and maintained in the exponential growth phase for 21 days. For subculturing, cells were trypsinized, counted and re-seeded with or without doxycycline. Cell counts from three biological replicates were used to calculate population doublings.

### RNA interference

For gene knockdown experiments, siRNAs were reverse transfected using Lipofectamine 2000 according to manufacturer’s instructions. Briefly, siRNA oligos were incubated with Lipofectamine 2000 and serum-free OptiMEM for 20 min at room temperature. Trypsinized cells were added directly into the siRNA/OptiMEM/Lipofectamine solution in the plate and were incubated as per individual experimental design. The siRNA sequences are listed in Supplementary Table 1.

### Clonogenic assay

Cells were seeded at a density of 2000 cells/well in triplicate in six-well plates. In the case of siRNA experiments, cells were transfected for 24 h before seeding to the plates. Growth medium was replenished every 3 days. For HPNE experiments, doxycycline was replenished every 3 days. HPNE cells and PDAC cells were allowed to grow for 6 days, except CAPAN-1 which was grown for 9 days. After 6-9 days depending on the cell lines, colonies were stained with 0.05% crystal violet in 1× PBS containing 1% methanol and 1% formaldehyde. The ImageJ plugin ColonyArea was used to automatically quantify stained colonies. The growth curves were generated by normalization to respective control group colony area. The SynergyFinder web application (Version 3.0) (https://synergyfinder.fimm.fi/) was used to generating synergy distribution heatmaps.

### Cell cycle and BrdU incorporation analysis

Growing cell monolayers were incubated with 10 µM BrdU for one hour. In experiments with adenovirus, cells were infected with adenoviruses at 1x10^10^ PFU/mL then cell monolayers were incubated with 10 µM BrdU for one hour. Cells were washed to remove unincorporated BrdU, trypsinized and suspended in fixing medium (65% PBS with 35% ethanol) overnight at 4 °C. Fixed cells were denatured using HCl and then neutralized with borax before stained with fluorescent anti-BrdU antibodies (FITC mouse anti-BrdU kit; 556028; BD) for half an hour. Nuclei were incubated in PBS containing 10 µg/mL of propidium iodide (PI) and 40 µg/mL of RNaseA for 1 h. Stained cells were analyzed by flow cytometry on an Accuri C6 flow cytometer (BD) using the manufacturer’s software. Replication licensing assay was performed as described previously (68). BrdU incopopration was calculated by subtracting median BrdU incorporation value for G1 phase cells from median BrdU incorporation value for S phase cells and normalized to control group.

### Gene expression and survival analysis

For RNA-seq, total RNA was isolated from hTERT-immortalized HPNE-DT cells with or without doxycycline induction of KRAS^G12V^ for six hours using QIAGEN RNeasy (QIAGEN, 74104). Libraries containing 250-300 bp cDNA inserts were polyA selected, prepared and sequenced on the Illumina platform PE150 by Novogene (Sacramento, CA). The TCGA pancreatic adenocarcinoma (TCGA-PAAD) data was retrieved using TCGAbiolinks R package (version 2.30.0). Samples defined as “high purity” (69) and “treatment-naïve” were used for gene expression analysis. Survival analysis and Kaplan-Meier curves were generated using the original TCGA_PAAD data and was performed using the following R packages: survival (version 3.2-13) and survminer (version 0.4.9).

### Mouse tumorigenesis studies

Animal experiments were approved by the Institutional Animal Care and Use Committees at UNC and performed according to guidelines. Suspensions of Pa03c, Pa02c, HPNE EV and KRAS^G12V^ cells at a concentration of 2.5x10^5^ cells in 40 µL sterile Matrigel-PBS were injected into the tail of the pancreas of three mice per group with the needle in a direction towards the head of the pancreas using a 0.30cc syringe and 30G needle. The needles were held still for 15-20s to prevent potential leaking. The pancreas was gently placed back into the body cavity and the surgery wound was sutured in layers, clipped and iodine will be applied to prevent infection. 0.1 mg/kg buprenorphine (Buprenex Injection, NDC 12496-07575) was injected subcutaneously into each mouse again at 24 hours. After the surgery, mice were placed on a heating pad for 5 mins to aid in recovery before returning to cage. Mice were monitored daily for 3 consecutive days after surgery and if required a third dose of buprenorphine was administered. Mice were monitored at least twice weekly. Animals were maintained for up to 6 weeks or until they needed to be euthanized based on humane endpoint criteria. Mouse KRAS^G12V^ induction was carried out by doxycycline-containing water (2 mg/mL, 5% w/v sucrose). Tumor growth was measured weekly using bioluminescence imaging (Revvity, IVIS Lumina III Series and XenoLight D-Luciferin – K+ Salt Substrate) performed by the UNC Animal Studies Core (see ‘Acknowledgements’). Pancreatic tumors were flash frozen and stored at -80C. For pathological analysis, tumors were added to molds pre-layered with O.C.T medium (Tissue-Tek, 62550) for cryofreezing. Cryofrozen OCT blocks were sectioned into 10 μm thick slices onto Superfrost plus slides (Fisherbrand, 1255015) and stored at -20°C. Slides were stained with Hematoxylin and eosin and images were taken at 10x and 2x resolution using a Zeiss Microscope.

### Iterative indirect immunofluorescence imaging

hTERT-HPNE EV and KRASG12V cells were seeded in glass-bottom 96-well plate treated with poly-L-lysine at 5,000 cells/well. Dox-induction of KRASG12V was done at the time of seeding at 200 ng/mL. After 3 days of induction, cells were pulse-labeled with EdU (10 µM) for 30 minutes, fixed with 4% PFA for 30 minutes then washed three times with DPBS (Corning, 21-031-CV). The blocking, imaging and elution iterations were done as described in (70). Briefly, each sample was incubated with 4i blocking solution (100 mM maleimide, 100 mM NH4Cl, and 1% BSA in PBS) for 1 hour, followed by incubation with diluted primary antibodies overnight at 4 °C. After 3 PBS washes, each sample was incubated with secondary antibodies. Samples were imaged on Nikon Ti Eclipse microscope. Stitched 8×8 images were acquired for each sample using the following filter cubes (Chroma): DAPI(383–408/425/435–485nm), GFP(450–490/495/500–550nm), Cy3(530–560/570/573–648nm), Cy5(590–650/660/663–738nm). After imaging each protein, samples were washed 3 times with ddH2O then antibodies were eluted and sequentially incubated with the next set of primary antibodies. The antibodies used are as follows: HA (Bethyl, A190-138A); phosphor-ERK (Cell Signaling Technology, 9101); phosphor-H2AX (Cell Signaling Technology, 80312); RB (Cell Signaling Technology, 9309); phosphor-RB (Cell Signaling Technology, 8516); phospho-p53 (S15) (Cell Signaling Technology, 9286); phospho-p21 (T145) (Abcam, ab47300); phospho- p27 (T157) (Abcam, ab805047); and phosphor-CHK1 (Cell Signaling Technology, 12302).

### PHATE analysis and visualization

The Potential of Heat-diffusion for Affinity-based Transition Embedding (PHATE) method (31796933) was applied using the previously mentioned feature set as input variables (we need to confirm the table below from Sam). The PHATE coordinates, which are data projections, position each cell relative to others based on their molecular signatures as determined by 4i protein markers. PHATE was run on z-normalized and filtered features with specific parameters for cell cycle maps: (Fig 1H: k-nearest neighbor (knn) = 200, gamma = 1, n=2). The proliferative cell cycle phases (G1/S/G2/M) in HPNE cells were manually annotated using distinct changes in DNA content. Data was visualized using Python (v3.11.8), numpy (1.26.4), pandas (2.0.3), phate (1.0.11) and Jupyter Notebooks (v2024.2.0). Statistical analysis was performed using one-way ANOVA with Tukey’s post- hoc test.

**Fig. 1.**
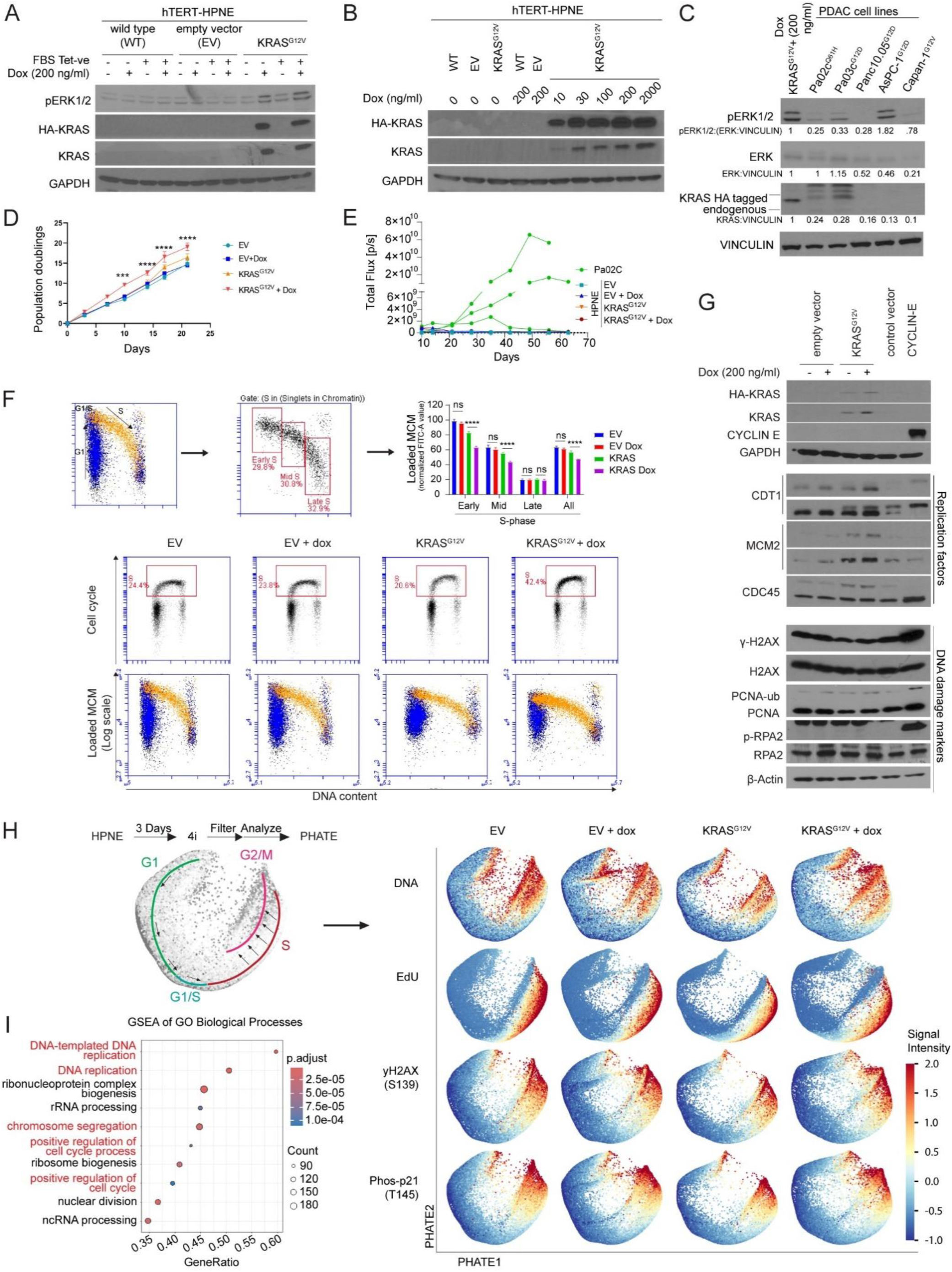
Establishing an HPNE cell system for modeling responses to KRAS-induced DNA replication stress (A)-(B) Immunoblots showing Doxycycline-inducible expression of HA-KRAS^G12V^ and p42/p44 MAPK phosphorylation in HPNE using regular and tetracycline-negative (Tet -ve) FBS. (C) Immunoblot showing Dox-inducible p42/p44 MAPK phosphorylation in HPNE and PDAC cell lines. (D) Growth curve showing Dox- inducible KRAS^G12V^ expression effect on proliferation. Error bars represent standard deviation (SD) for biological triplicates. Statistical analysis: two-way ANOVA followed by Tukey’s multiple comparisons test, * = *p* < 0.05, ** = *p* < 0.01, *** = *p* < 0.001, **** = *p* < 0.0001, ns = not significant. (E) Plot showing tumor growth of orthotopically implanted HPNE and PDAC cell lines into the pancreas of nude mice (n = three mice per group). (F) Flow cytometry profiles showing the effect of Dox-inducible KRAS^G12V^ expression on BrdU incorporation (upper panels) and chromatin-loading of MCM2 (middle panels). The bar charts show quantification of chromatin-loaded MCM (normalized FITC-A value) under the different experimental conditions. Error bars represent standard deviation (SD) for biological triplicates. Statistical analysis: two-way ANOVA followed by Tukey’s multiple comparisons test, * = *p* < 0.05, ** = *p* < 0.01, *** = *p* < 0.001, **** = *p* < 0.0001, ns = not significant. (G) Effects of Dox-inducible KRAS^G12V^ or adenovirus-transduced Cyclin E expression on the indicated DNA damage markers and DNA replication factors. HPNE cells were infected with Adenovirus expressing CyclinE1 (or ’empty’ control virus) at 1x10^10^ PFU/mL (H) PHATE plots indicating cell cycle trajectories and DNA damage markers of empty vector and KRAS^G12V^ -expressing HPNE and their relevant empty vector controls after 3 days of Dox-treatment. (I) GO analysis of KRAS^G12V^-induced mRNAs compared to empty vector in HPNE showing top 10 GO terms.

### DNA fiber analysis of DNA replication dynamics

Growing cells were labeled with 25 µM chloro-deoxyuridine (CIdU; C6891; Sigma-Aldrich) for 20 min. Cells were then washed twice with warm medium and immediately labeled with 250 µM iodo-deoxyuridine (IdU; I7125; Sigma-Aldrich) for 20 min. Labeled cells were harvested, washed in PBS, and added to glass slides. Cells were lysed in spreading buffer (200 mM Tris-HCl, pH 7.5, 50 mM EDTA, 0.5% SDS), and DNA fibers were spread across the slide by tilting the slides at an angle. Air dried DNA fiber slides were fixed in a 3:1 mix of methanol and acetic acid for 10 min. DNA fibers were denatured in 2.5M HCL for 1h, incubated in blocking solution (1% BSA, 1% Tween 20 in PBS) for 30 min, and then incubated with mouse anti-BrdU (BD Biosciences, 347583; for IdU detection) and rat anti-BrdU (Abcam, A6326; for CldU detection) primary antibodies for 1h in the dark. Subsequently, DNA fibers were incubated with secondary antibodies, anti-mouse Alexaflour 488 and anti-rat Alexaflour 555. Flourescently-labelled DNA fibers were imaged using an Andor Dragonfly Spinning Disk Confocal Microscope (OXFORD Instruments, England) mounted on a Leica DMi8 microscope stand, equipped with an HC PL APO 100×/1.40 OIL CS2 Leica objective. The lengths of CldU (AF 555; red)- and IdU (AF 488; green)-labeled fibers were measured using the ImageJ software, and micrometer values were converted into kilobases using the conversion factor 1 µM = 2.94 kb (one bp corresponds to ∼340 pm). A minimum of 50 representative DNA fibers were counted for each experimental condition.

### Live cell microscopy

To generate hTERT-HPNE cell lines stably expressing GFP-H2B, EV and KRAS^G12V^-HA cells were infected with lentivirus expressing GFP-H2B and selected by growth in medium containing hygromycin B (ThermoFisher). Cell lines with stable GFP-H2B expression were seeded on Chambered Coverglass from Lab-Tek II (ThermoFisher, 155382). Cell lines were treated with doxycycline and transfected with siRNA 24 h before imaging. Time-lapse microscopy was performed on a Keyence BZ-X810 using a 20× objective. Images were taken at 4 min interval for 24 h. Best focus projections of the time series were exported into AVI format. Image sequences were generated using ImageJ and manually quantified.

### Immunofluorescence (IF)

Two days post siRNA transfection and doxycycline treatment, HPNE cells were washed twice with PBS, fixed with 2% paraformaldehyde (PFA) for 15 minutes, washed twice with PBS, permeabilized with 0.2% triton-X 100 for 5 minutes, washed three times with PBS, and blocked with 3% bovine serum albumin (BSA). This was followed by primary antibody (53BP1, sc-22760, 1:200 dilution) incubation for 1h, three PBS washes, secondary antibody incubation for 1h, and three washes in PBS. Finally, coverslips were mounted on the slides using fluoroshield with 4’,6-diamidino-2-phenylindole (DAPI) (#H-1200 Vector laboratories).

For analysis of 53BP1 foci, high resolution images were taken using an Andor Dragonfly Spinning Disk Confocal Microscope (OXFORD instruments, England) mounted in a Leica Dmi8 microscope stand, equipped with an HC PL APO 100X/1.40 OIL CS2 Leica objective. The pinhole size was set to 40 μm, and the camera used was a Zyla Plus 4.2MP sCMOS, featuring a resolution of 2408 x 2048 pixels and an effective pixel size of 0.063 μm. Excitation lasers included a 405 nm laser for DAPI and a 488 nm laser for Alexa Fluor 488. Emission filters used were 445/46 for DAPI and 521/38 for Alexa Fluor 488.

### Statistics and reproducibility

Statistical analyses were performed using Microsoft Excel and GraphPad Prism. The student’s t-test was used to determine P values for all data involving comparisons between two groups. Results are expressed as the mean ± standard error of the mean (SEM) or standard deviation (SD) of two or more independent experiments as indicated in the legends. The *P* values are indicated in the Figure legends. All biological and biochemical experiments were performed with appropriate internal negative and/or positive controls as indicated.

## RESULTS

### Establishing an experimental system for defining responses to oncogenic KRAS in normal pancreatic epithelial cells

To model PDAC-relevant oncogenic events we engineered Human Pancreatic Nestin-Expressing cells (HPNE, TERT-immortalized normal epithelial cell PDAC precursors) to express oncogenic KRAS^G12V^ in a Doxycycline (Dox)-inducible manner (Fig. 1A). As expected, Dox-induction of KRAS^G12V^ expression led to increased phosphorylation of ERK1/2, downstream targets of the KRAS pathway. A Dox dose of 200 ng/ml induced levels of KRAS^G12V^ and pERK comparable to those found in pancreatic adenocarcinoma (PDAC) cell lines (Fig. 1B and Fig. 1C). This level of KRAS^G12V^ expression led to increased cell proliferation rates *in vitro* (Fig. 1D). However, when orthotopically implanted into the pancreas of nude mice, KRAS^G12V^-expressing HPNE-cells did not form tumors. In contrast, orthotopically-implanted PDAC cell lines readily induced tumor growth (Fig. 1E and Supplementary Figure S1B-F). Therefore, Dox-induced KRAS^G12V^ expression in HPNE is a model for pre-neoplasia. In flow cytometry experiments, Dox induction of KRAS^G12V^ stimulated cells to enter S-phase with reduced levels of minichromosome maintenance (MCM)- loaded chromatin (Fig. 1F), indicating that oncogenic KRAS accelerates G1 at the expense of DNA replication licensing in this system. In contrast with overexpressed Cyclin E (a commonly-used oncogene for modeling DNA replication stress), the levels of Dox-induced KRAS^G12V^ expression attained in HPNE did not elicit high levels of DNA replication stress as shown by flow cytometry analysis of BrdU-labelled cells (Fig. 1F and Supplementary Figure S1A) and DNA damage markers such as γH2AX, PCNA monoubiquitylation (PCNA-ub) and pRPA as shown by western blot (Fig. 1G).

To ask whether higher resolution analysis would reveal KRAS^G12V^-induced cell cycle defects in HPNE we performed iterative indirect immunofluorescence imaging (4i) (70). As shown in Fig. 1H and Supplementary Figure S2, our single cell resolution 4i analyses confirmed that KRAS^G12V^ expression induced no major branch points or cell groups. Of the protein markers we examined, only levels of γΗ2ΑΧ (S139) in S-phase, phosphorylated-TP53 (S15) in G2/M-phases and phosphorylated-p27kip (T157) were modestly increased as a result of KRAS^G12V^ expression (Supplementary Figure S3). We conclude that low-levels of KRAS^G12V^ signaling promotes proliferation without significantly perturbating the cell cycle or inducing replication stress and DNA damage markers in HPNE.

Using RNASeq we confirmed that KRAS^G12V^ expression in HPNE induced a robust transcriptional program including expression of DNA replication and cell cycle genes (Fig. 1I). Taken together, our results suggest that the DNA replication stress induced by pathologically-relevant levels of KRAS^G12V^ signaling is tolerated by HPNE. Therefore, HPNE provides an excellent opportunity to define mechanisms that help tolerate KRAS^G12V^- induced replication stress.

### siRNA screening identifies *TRIP13* as a dependency of KRAS^G12V^-expressing HPNE cells

Expression of DNA repair genes is often upregulated according to the needs of cells. For example, Homologous Recombination (HR)-deficient cancers upregulate POLQ, a DNA polymerase that mediates a back-up DNA repair mechanism needed to tolerate HR-deficiency (71). By analogy, we hypothesized that cells expressing KRAS^G12V^ might upregulate genome maintenance factors involved in tolerating KRAS-induced DNA replication stress. Therefore, we used RNAseq data to identify DDR genes (supplementary table 3, gene list) that were differentially expressed (adj. p-value < 0.05; absolute (Log2FC) > 0.5) between control and KRAS^G12V^-expressing HPNE. As shown in Fig. 2A, KRAS^G12V^ expression resulted in 109 and 26 DDR genes being upregulated and downregulated, respectively. We also interrogated gene expression data for TCGA-PAAD and identified 43 and 17 DDR genes being upregulated and downregulated in primary KRAS-mutated tumors when compared with normal pancreas (Fig. 2B). The heatmaps in Fig. 2C and Fig. 2D show the relative expression levels of DDR genes that were upregulated in KRAS^G12V^-expressing HPNE cells (when compared with parental HPNE) and KRAS-mutant pancreatic cancers (when compared with normal pancreatic tissue). 15 of the DDR genes that were upregulated following KRAS^G12V^ induction in HPNE cells were also expressed at high levels in KRAS-mutant pancreatic tumors (Fig. 2E). The DDR genes that were upregulated in both mutant KRAS-expressing groups are highlighted in red in Fig. 2C and Fig. 2D. We generated a library of siRNAs targeting 17 of the DDR genes that were upregulated in KRAS^G12V^-expressing HPNE (ARID1A, BRCA2, CDCA7L, FANCI, HELLS, MCM6, RAD54B, RNF168), in mutant KRAS-expressing pancreatic tumors (MNAT1, TK1, SMYD3), or in both groups (AURKA, CDC6, MELK, PLK1, RAD51, TRIP13). Using the resulting siRNA library (which targets KRAS-inducible and PDAC-associated DDR mRNAs), we screened for genes that were required for cell viability and ongoing DNA synthesis only in KRAS-active HPNE. The experimental workflow for our screen is depicted in Fig. 3A. Fig. 3B summarizes our screen results and provides an aerial view of how the DDR-directed siRNAs in our library affected survival and on-going DNA synthesis of EV-HPNE and KRAS^G12V^-HPNE +/- Dox.

**Fig. 2.**
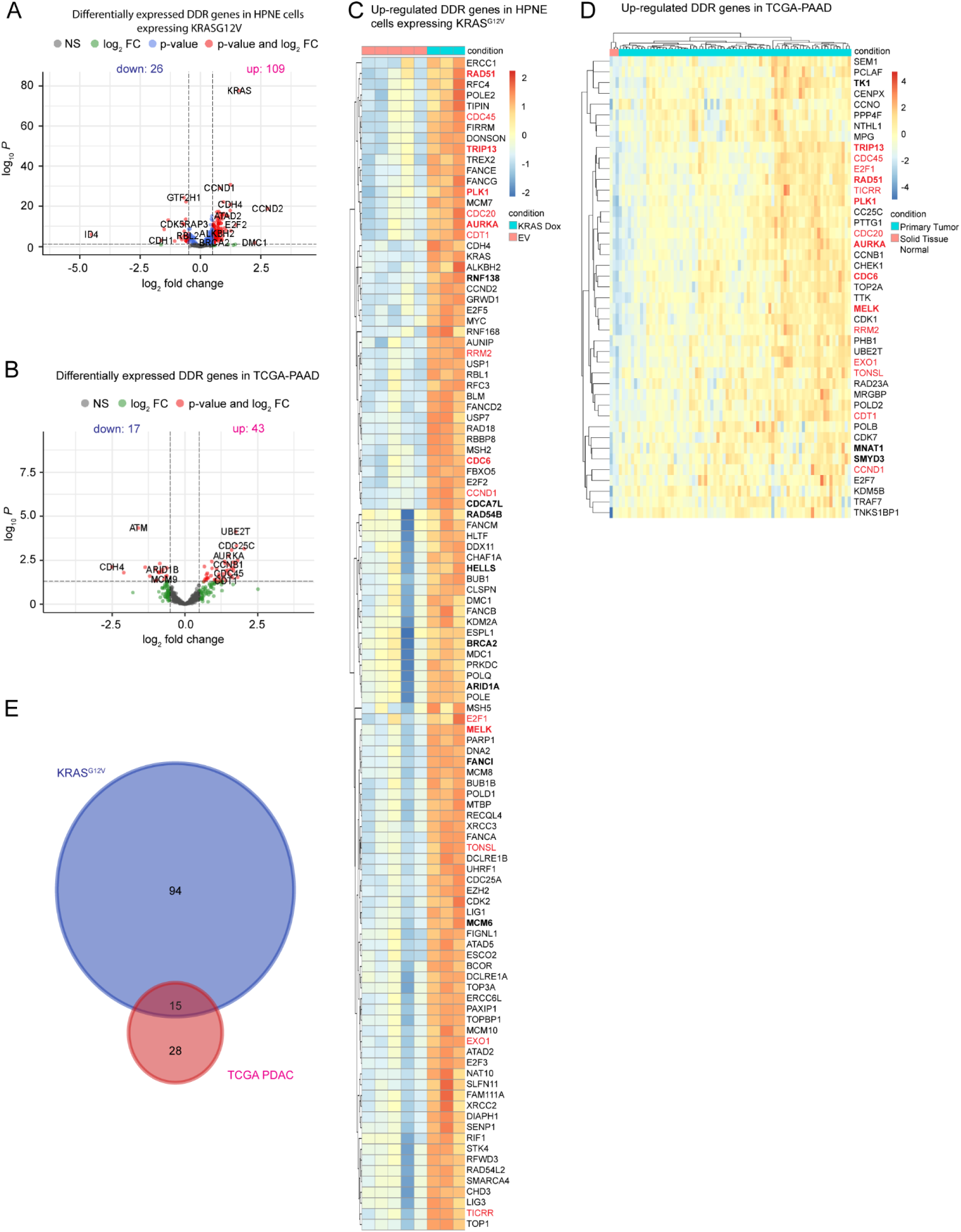
Analysis of the HPNE transcriptome reveals KRAS-inducible DDR genes (A) Volcano plot showing log2 fold changes (FC) in mRNA expression of DDR genes against -log10 adjusted p-value for hTERT-HPNE expressing KRAS^G12V^ against control with wild type KRAS. (B) Volcano plot showing log2 fold changes (FC) in mRNA expression of DDR genes against -log10 adjusted p-value TCGA-PAAD primary tumors against solid tissue normal pancreas. (C) Heatmap showing up regulated DDR genes in hTERT-HPNE expressing KRAS^G12V^ against control with wild type KRAS. (D) Heatmap showing up regulated DDR genes in TCGA-PAAD primary tumors against solid tissue normal pancreas. (E) Venn diagram showing the numbers of upregulated DDR genes in hTERT-HPNE expressing KRAS^G12V^, TCGA-PAAD primary tumors, and overlap between them. (BOLD and black font represents 15 upregulated DDR genes common between hTERT-HPNE expressing KRAS^G12V^ and TCGA-PAAD primary tumors; BOLD and red fonts represent the genes selected for siRNA screening).

**Fig. 3.**
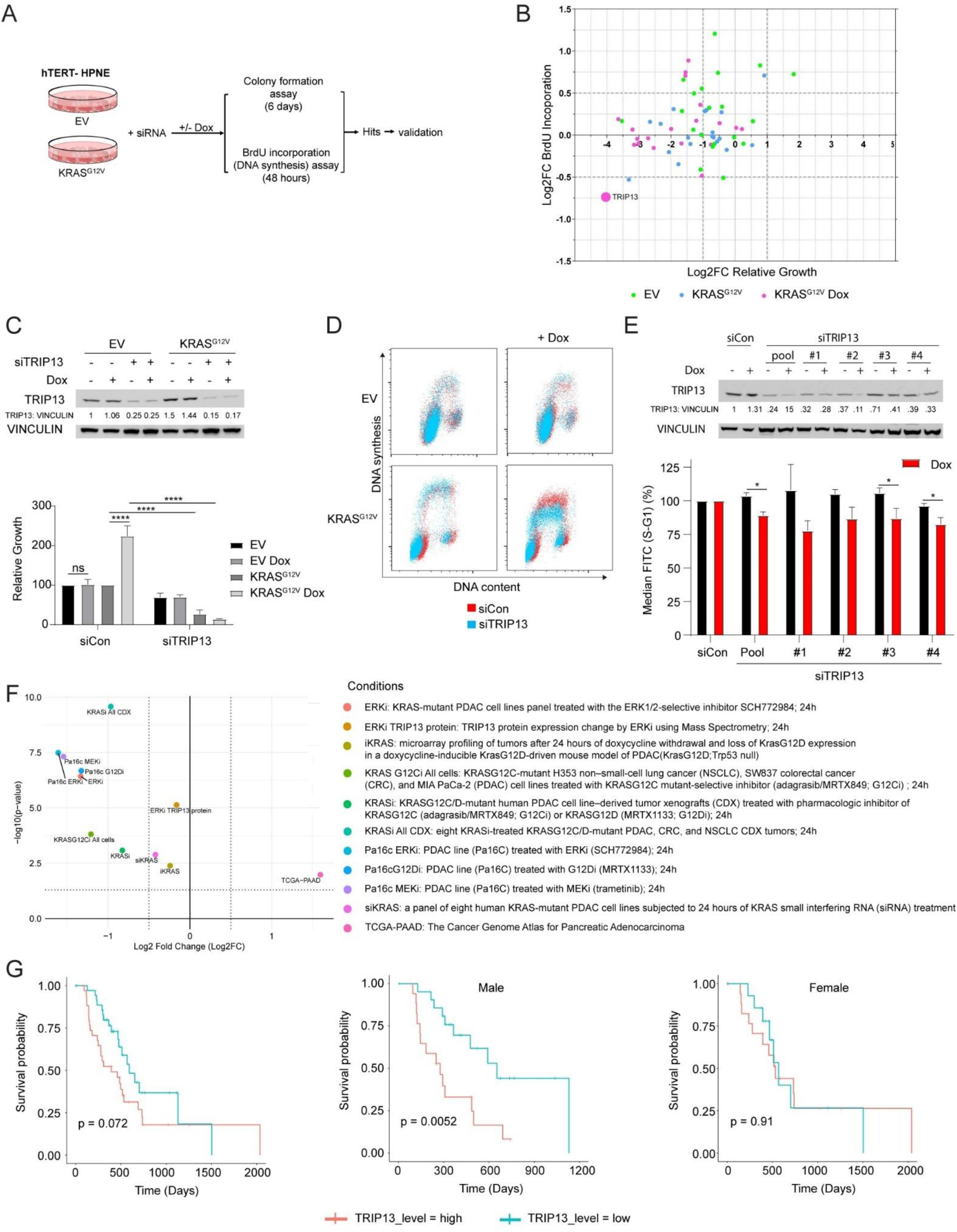
siRNA Screening identifies TRIP13 as a dependency of KRAS-expressing HPNE. (A) Experimental workflow of siRNA screen for DDR-dependencies of KRAS^G12V^-expressing HPNE. (B) Plot showing effect of library siRNAs on clonogenic survival and BrdU incorporation of empty vector and KRAS^G12V^- expressing HPNE. (C) Immunoblot validating siRNA-mediated knockdown of TRIP13 (upper panel) and bar chart showing effect of TRIP13 knockdown on clonogenic survival in empty vector and KRAS^G12V^-expressing HPNE. Error bars represent standard deviation (SD) for biological triplicates. Statistical analysis: two-way ANOVA followed by Tukey’s multiple comparisons test, * = *p* < 0.05, ** = *p* < 0.01, *** = *p* < 0.001, **** = *p* < 0.0001, ns = not significant. (D) Flow cytometry plots showing effect of TRIP13 siRNA on cell cycle profiles and BrdU incorporation in empty vector and KRAS^G12V^-expressing HPNE. (E) Results of immunoblot (upper panel) and flow cytometry analyses (lower panel) showing effect of multiple independent TRIP13-directed siRNAs on BrdU incorporation in KRAS^G12V^-expressing HPNE. Error bars represent standard deviation (SD) for biological triplicates. Statistical analysis: Student’s *t*-test, * = *p* < 0.05, ** = *p* < 0.01, *** = *p* < 0.001, **** = *p* < 0.0001, ns = not significant. (F) Volcano plot showing effects of KRAS and ERK status on TRIP13 expression. (G) Kaplan-Meier curves showing overall survival of TRIP13 expression in TCGA-PAAD.

Our screening experiments identified *TRIP13* as a requirement for both clonogenic survival and ongoing DNA synthesis in KRAS^G12V^-expressing HPNE but not in control HPNE lacking mutant KRAS (Supplementary Figures S4A, S4B and S4C). In secondary validation experiments, KRAS^G12V^ expression led to more than two-fold increase in clonogenic survival in HPNE treated with a control siRNA (Fig. 3C). However, in TRIP13- ablated cells, KRAS induction led to around five-fold decrease in clonogenic survival (Fig. 3C). Our flow cytometry analyses showed that TRIP13-depletion led to reduced DNA synthesis rates only in KRAS^G12V^- expressing HPNE (Fig. 3D). We validated the role of TRIP13 in averting DNA replication stress using four independent siRNAs targeting TRIP13 (Fig. 3E). We conclude that TRIP13 is specifically required for DNA synthesis and viability of HPNE expressing oncogenic KRAS. We asked whether TRIP13 is also important for tolerating replication stress induced by other oncogenes. As shown in Supplementary figure S5A-D, TRIP13 depletion also led to reduced DNA synthesis and decreased cell viability in HPNE cells in which we ectopically expressed Cyclin E or MYC oncogenes. Based on our analysis with representative important human oncogenes (KRAS, Cyclin E, MYC), we suggest that TRIP13 may have broad roles in sustaining DNA synthesis and viability of oncogene-expressing cells.

Next we determined the relationship between TRIP13 expression and KRAS-ERK signaling in cancer. Klomp et al. recently performed transcriptional profiling of a large panel of cell lines and PDX models in which KRAS/ERK signaling was conditionally ablated to define a KRAS/ERK-dependent gene signature (72). We interrogated the transcriptome data generated by Klomp et al. to test relationships between TRIP13 expression and KRAS/ERK signaling. As shown in Fig. 3F, conditional ablation of KRAS/ERK signaling was associated with reduced *TRIP13* expression in all datasets. In a TCGA cohort of PDAC patients, high TRIP13 expression (upper quartile) was associated with reduced survival probability (Fig. 3F). Interestingly, the correlation between high TRIP13 expression and reduced survival was evident only in male patients (Fig. 3F, Supplementary Figure 5E). These clinical data are also potentially consistent with a role for TRIP13 in sustaining tumors harboring oncogenic KRAS.

### TRIP13 sustains DNA replication in KRAS-expressing cells via the HR pathway

The reduced DNA synthesis rates that we observed in TRIP13-ablated cells expressing oncogenic KRAS could potentially be due to changes in replication fork velocity or altered origin firing. To distinguish between these mechanisms, we investigated the role of TRIP13 in DNA replication dynamics using DNA fiber analysis. As shown in Fig. 4A TRIP13-depletion led to reduced DNA replication fork speeds specifically in KRAS^G12V^-expressing cells. Loss of TRIP13 in KRAS^G12V^-expressing cells also led to an increase in numbers of stalled forks (Supplementary Figure S6), but did not significantly reduce new origin firing.

**Fig. 4.**
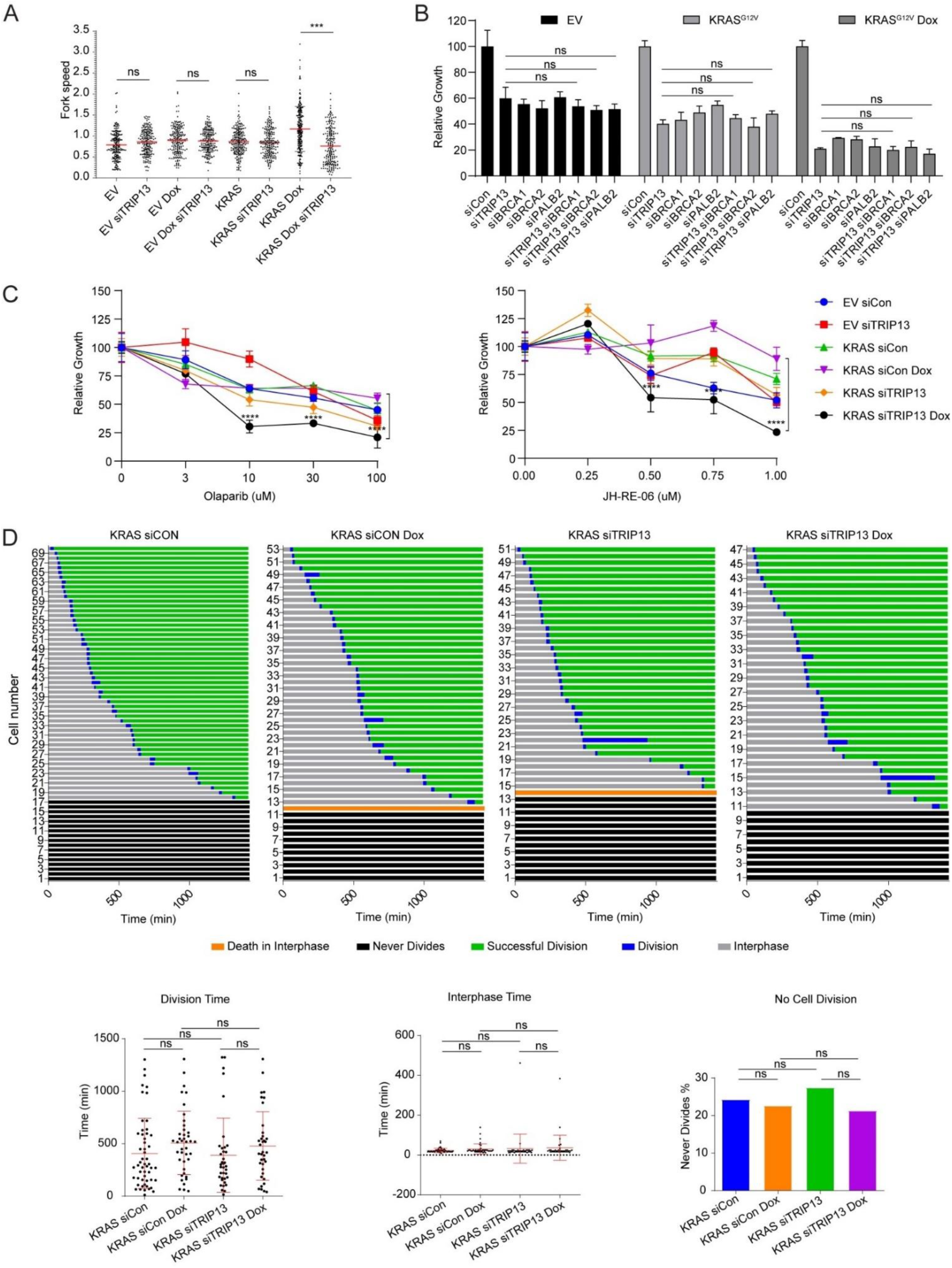
TRIP13 is epistatic with HR genes for tolerance of KRAS-induced DNA replication stress in HPNE cells. (A) Results of DNA fiber assays showing effects of KRAS^G12V^ expression and TRIP13 knockdown on DNA replication fork velocities in HPNE. Error bars represent standard deviation (SD) for biological triplicates. Statistical analysis: one-way ANOVA followed by Tukey’s multiple comparisons test, * = *p* < 0.05, ** = *p* < 0.01, *** = *p* < 0.001, **** = *p* < 0.0001, ns = not significant. (B) Bar chart showing effects of siRNAs against TRIP13, BRCA1, BRCA2, and PALB2 on colony survival assays in empty vector and KRAS^G12V^-expressing HPNE. Error bars represent standard deviation (SD) for triplicate. Statistical analysis: one-way ANOVA followed by Tukey’s multiple comparisons test, * = *p* < 0.05, ** = *p* < 0.01, *** = *p* < 0.001, **** = *p* < 0.0001, ns = not significant. (C) Results of clonogenic survival assays showing dose-dependent effects of olaparib (left panel) and JH-RE-06 (right panel) on viability of empty vector and KRAS^G12V^-expressing HPNE. Error bars represent standard deviation (SD) for triplicate. Statistical analysis: two-way ANOVA followed by Tukey’s multiple comparisons test, * = *p* < 0.05, ** = *p* < 0.01, *** = *p* < 0.001, **** = *p* < 0.0001, ns = not significant. (D) Results of live cell imaging showing mitotic fates of control and Dox-induced KRAS^G12V^-HPNE cells. Error bars represent standard deviation (SD) for biological triplicates. Statistical analysis: two-way ANOVA followed by Tukey’s multiple comparisons test, * = *p* < 0.05, ** = *p* < 0.01, *** = *p* < 0.001, **** = *p* < 0.0001, ns = not significant.

Next, we considered the potential mechanisms by which TRIP13 facilitates replication fork movement. TRIP13 is an AAA+ ATPase that regulates the cell cycle and genome maintenance factors MAD2 and REV7 (73).

Mechanistically, TRIP13 converts MAD2 and REV7 from ‘closed’ to ‘open’ conformations dissociating them from their binding partners. TRIP13-mediated dissociation of MAD2 and REV7 complexes inhibits Trans- Lesion Synthesis (TLS, a damage-tolerant mode of DNA synthesis), promotes Homologous Recombination (HR) and relieves inhibition of mitotic progression. TRIP13 ablation is expected to promote Polζ-mediated TLS, inhibit HR, and relieve APC inhibition to promote mitotic progression. Therefore, we systematically tested whether increased TLS, reduced HR, or changes in mitotic exit accounted for the TRIP13-dependency of KRAS^G12V^-expressing cells.

First, we performed epistasis analysis to investigate the relationship between TRIP13 and HR factors in KRAS^G12V^-expressing cells. We used siRNA to knock down key factors in the HR pathway (BRCA1, BRCA2, and PALB2) individually or in combination with TRIP13, then determined the impact of these ablations on clonogenic survival of control or KRAS^G12V^-expressing cells. As shown in Fig. 4B, knockdown of BRCA1, BRCA2 or PALB2 phenocopied the survival defects of TRIP13-depleted cells. Moreover, co-depletion of TRIP13 with BRCA1, BRCA2, or PALB2 did not lead to additive effects on colony survival. The results of Fig. 4B suggest that TRIP13 participates in the same pathway as BRCA1, BRCA2, or PALB2 to maintain HR and promote survival of KRAS^G12V^-expressing cells. HR-compromised cells are typically sensitive to PARP inhibitors (74). As shown in Fig. 4C, TRIP13-depleted cells were sensitized to the PARP inhibitor Olaparib in a KRAS^G12V^-dependent manner, further indicating that TRIP13 sustains KRAS^G12V^-expressing cells via an HR-mediated mechanism.

Interestingly, TRIP13 knockdown also led to reduced numbers of nuclear 53BP1 foci across all of our HPNE cell lines (Supplementary Figure S7A), suggesting that TRIP13 may also have a role in promoting Non-Homologous End Joining (NHEJ), as also suggested by other investigators (75,76). However, based on the epistatic relationship between TRIP13 and HR genes (Fig. 4B), we infer that the role of TRIP13 in HR is most important for promoting tolerance of KRASG12V-induced replication stress.

Next, we tested if the effect of TRIP13-depletion on survival of KRAS^G12V^-expressing cells was mediated via TLS. We reasoned that if excessive TLS due to accumulation of REV7-REV3 complexes accounts for the survival defects of TRIP13-depleted (KRAS^G12V^-expressing) cells, these defects should be rescued by TLS inhibition. Therefore, we used the REV1 inhibitor JH-RE-06 (77) to inhibit TLS in KRAS^G12V^-expressing HPNE cells that were conditionally treated with TRIP13 siRNA. Interestingly, TLS inhibition did not rescue the reduced viability of TRIP13-depleted KRAS^G12V^-expressing cells (Fig. 4C). On the contrary, JH-RE-06 treatment further reduced viability of TRIP13-deficient cells expressing oncogenic KRAS (Fig. 4C). We conclude that the sensitivity of TRIP13-depleted cells to oncogenic KRAS is not caused by excessive TLS. Instead, we infer that the HR-deficiency of TRIP13-depleted cells leads to increased dependency on TLS to facilitate tolerance of KRAS^G12V^-induced DNA replication stress.

Finally, to determine if the survival defects of TRIP13-depleted cells expressing KRASG12V were mediated via dysregulation of the mitotic MAD2B-APC signaling axis we directly measured mitotic progression using live cell imaging. We generated HPNE and HPNE-KRAS^G12V^ cells stably expressing a GFP-histone construct, treated the resulting cells conditionally with TRIP13 siRNA and visualized chromosome dynamics over a 24 h period. Movies were analyzed and fate maps were generated for all experimental conditions. As shown in Fig. 4D, TRIP13 depletion in KRAS^G12V^-expressing cells did not affect mitotic timing or post-mitotic fates of dividing cells. We conclude that TRIP13 does not significantly affect MAD2/APC function or mitotic exit in KRAS^G12V^- expressing cells. Instead TRIP13 promotes survival of KRAS^G12V^-expressing cells by maintaining HR.

### TRIP13 is a dependency of established PDAC cells

Because our experiments with a pre-neoplasia model showed that TRIP13 sustains KRAS^G12V^-expressing cells, we next asked whether TRIP13 was also necessary for survival of established PDAC cells expressing oncogenic KRAS. Therefore, we investigated the TRIP13- dependency of a panel of five established PDAC cell lines, including homologous recombination-deficient (HRD) CAPAN-1 cells. As shown in Fig. 5A and Fig. 5B, treatment with TRIP13 siRNA suppressed clonogenic survival and DNA synthesis of HR-sufficient PDAC cell lines (AsPc-1, Panc10.05, Pa02C, and Pa03C).

**Fig. 5.**
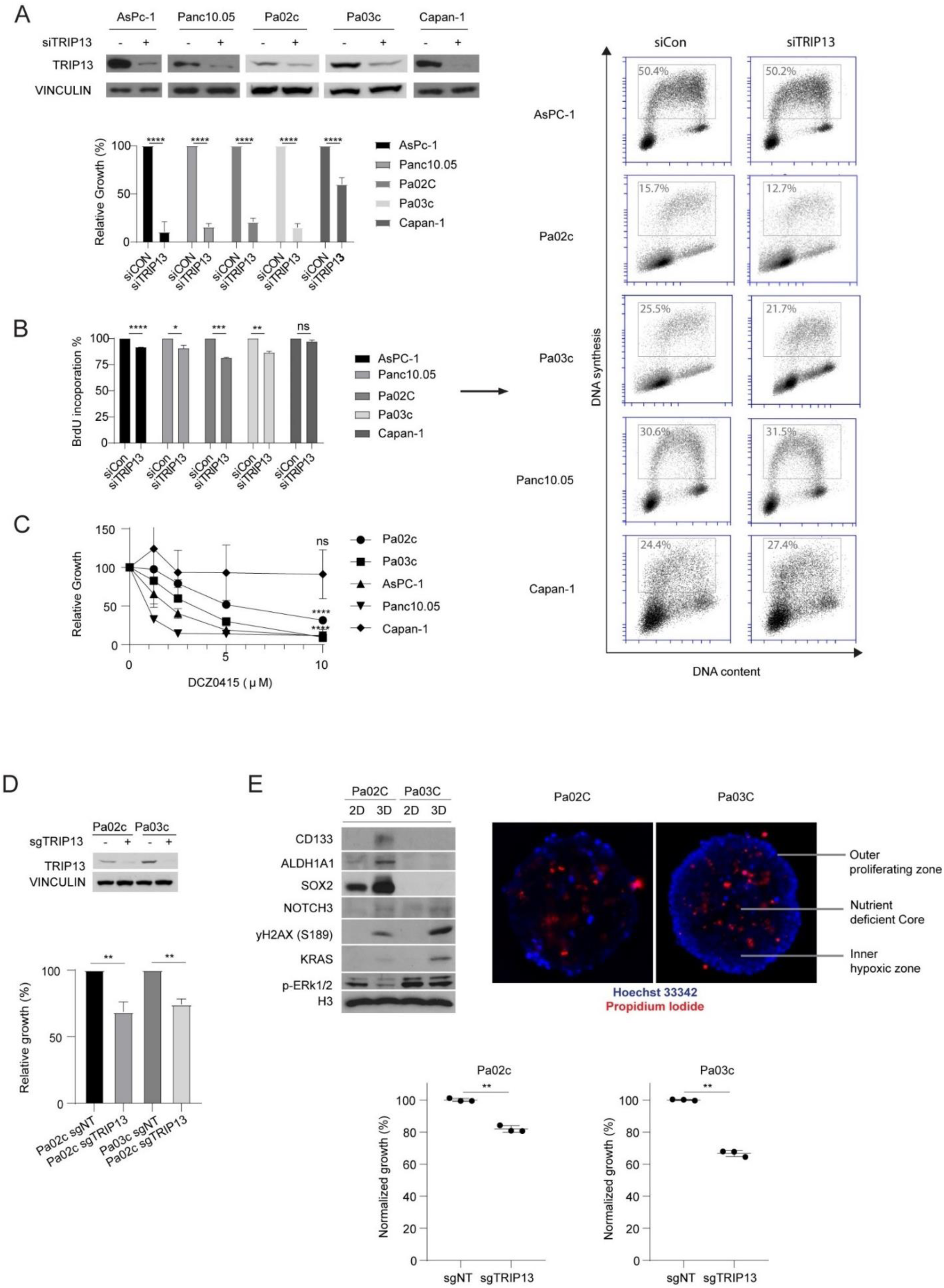
PDAC cell lines require TRIP13 for survival and DNA synthesis. (A) Immunoblots validating TRIP13 knockdown using siRNA in PDAC cell lines (upper panel) and effects of TRIP13 siRNA on clonogenic survival of PDAC cells (lower panel). Error bars represent standard deviation (SD) for triplicate. Statistical analysis: Student’s *t*-test, * = *p* < 0.05, ** = *p* < 0.01, *** = *p* < 0.001, **** = *p* < 0.0001, ns = not significant. (B) Effect of TRIP13 siRNA on cell cycle distribution and BrdU incorporation in PDAC cell lines. Error bars represent standard deviation (SD) for triplicate. Statistical analysis: two-way ANOVA followed by Tukey’s multiple comparisons test, * = *p* < 0.05, ** = *p* < 0.01, *** = *p* < 0.001, **** = *p* < 0.0001, ns = not significant. (C) Dose dependent inhibition of clonogenic survival by DCZ0415 in PDAC cell lines. Error bars represent standard deviation (SD) for triplicate. Statistical analysis: Student’s *t*-test, * = *p* < 0.05, ** = *p* < 0.01, *** = *p* < 0.001, **** = *p* < 0.0001, ns = not significant. (D) Immunoblots validating TRIP13 knockdown using siTRIP13 in PDAC cell lines (upper panel) and effects of sgTRIP13 on clonogenic survival of PDAC cells (lower panel). Error bars represent standard deviation (SD) for triplicate experimental samples. Statistical analysis: Student’s *t*-test, * = *p* < 0.05, ** = *p* < 0.01, *** = *p* < 0.001, **** = *p* < 0.0001, ns = not significant. (E) The upper panels show immunofluorescence microscopy images of 3D spheroids of PDAC cell lines (Blue = Hoechst 33342 and Red = Propidium iodide) and immunoblots comparing expression levels of the indicated proteins between 2D and 3D cultures. The lower panel shows the effect of sgTRIP13 treatment on the growth of PDAC cells in spheroid culture as determined using a GFP competitive growth assay.

However, clonogenic survival and DNA synthesis of BRCA2-deficient CAPAN-1 cells were relatively insensitive to TRIP13-depletion (Fig. 5A and Fig. 5B).

In the HR-sufficient PDAC cell lines clonogenic survival was reduced after treatment with the TRIP13 inhibitor DCZ0415 (Fig. 5C). However, clonogenic survival of HR-deficient CAPAN-1 cells was relatively resistant to pharmacological inhibition of TRIP13 (Fig. 5C). Similar to our results with DCZ0415-treatment, genetic ablation of *TRIP13* (using sgRNA) led to reduced proliferation of Pa02c and Pa03c PDAC cell lines (Fig. 5D). We recently reported that cancer cells in 3D culture (which recapitulate hallmark characteristics of tumors more closely than cells in monolayer culture) often have different requirements for stress tolerance (78). Therefore, we established conditions for growing Pa02c and Pa03c cells in 3D culture. As shown in Fig. 5E, both PDAC cell lines seeded in ultra-low attachment plates organized into 3D spheroid structures with radial gradients of proliferation and death that mimic how cells organize in tumors. As expected from our previous work (78), immunoblotting of cell extracts revealed differences in expression of cancer stemness markers between 2D and 3D cultures of PDAC cells (Fig. 5E). Interestingly, in both Pa02c and Pa03c cell lines, KRAS expression was elevated during 3D growth when compared with monolayers. Consistent with the results of our previous experiments using monolayer cultures, *TRIP13* ablation (using sgRNA) also led to reduced viability of Pa02c and Pa03c cells in 3D culture (Fig. 5E). Increasingly spheroids and ‘tumorsphere’ culture is being adopted as a preclinical model. The results of Fig. 5E strengthen the notion that TRIP13 is important for PDAC cells to grow as organized and pathologically-relevant 3D structures.

### TRIP13 confers resistance to therapeutic DNA damaging agents

Next, we asked whether *TRIP13* loss can also be leveraged as a vulnerability to enhance sensitivity to therapeutic agents. Similar to KRAS^G12V^- expressing HPNE cells, *TRIP13*^-/-^ Pa02c and Pa03c PDAC cell lines (Supplementary Figure S5B), displayed a hallmark *BRCA*-ness phenotype, namely PARPi-sensitivity (Fig. 6A). Recent work suggests that *BRCA*- deficiencies lead to accumulation of extensive single-stranded DNA in the genome (79). Therefore, we predicted that *TRIP13*-deficiency would lead to a dependency on TLS-mediated ssDNA gap-filling. Consistent with this prediction, *TRIP13*^-/-^ PDAC cells were sensitive to the TLS inhibitor JH-RE06 (Fig. 6A). *TRIP13*- deficiency also sensitized PDAC cells to gemcitabine (a first-line therapy for pancreatic cancer) and the WEE1- inhibitor AZD1775 (which has been explored as a potential treatment for PDAC) (Fig. 6A). The olaparib- and JH-RE-06-sensitivities of *TRIP13*-PDAC cell lines were evident when we performed these experiments using multiple platforms including clonogenic assays (Fig. 6A), CellTiter-Blue® Cell Viability Assays (Fig. 6B) and 3D spheroid cultures (Fig. 6C). Taken together these experiments demonstrate the robustness of the synergy between TRIP13-deficiency and TLS or PARP inhibition.

**Fig. 6.**
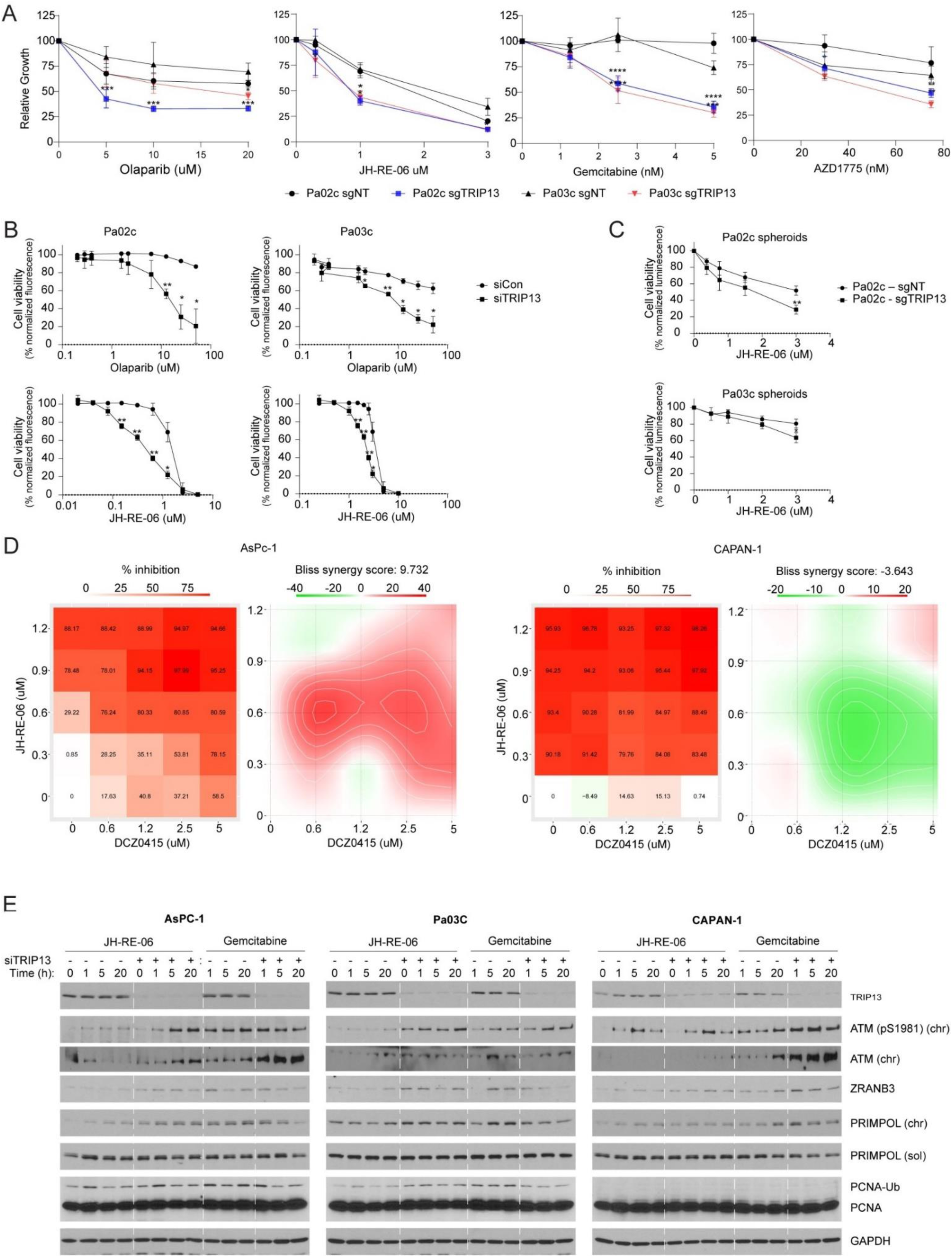
TRIP13-loss sensitizes PDAC cell lines to therapeutic agents (A) Dose-dependent effects of pharmacological agents (JH-RE-06, Gemcitabine, Olaparib and AZD1775) on clonogenic survival of Pa02C and Pa03C cells after conditional treatment with TRIP13 siRNA. Error bars represent standard deviation (SD) for triplicate. Statistical analysis: Student’s *t*-test, * = *p* < 0.05, ** = *p* < 0.01, *** = *p* < 0.001, **** = *p* < 0.0001, ns = not significant. (B) Results of CellTiter-Blue® assays showing dose-dependent effects of pharmacological agents (JH-RE-06 and Olaparib) on viability of Pa02C and Pa03C cells after conditional treatment with TRIP13 siRNA. Error bars represent standard deviation (SD) for biological duplicates. Statistical analysis: *t*-test, * = *p* < 0.05, ** = *p* < 0.01, *** = *p* < 0.001, **** = *p* < 0.0001, ns = not significant. (C) Results of CellTiter-Blue® assays showing dose-dependent effects of JH-RE-06 on viability of Pa02C and Pa03C 3D cells in spheroid culture after conditional treatment with sgTRIP13. Error bars represent standard deviation (SD) for biological duplicates. Statistical analysis: *t*-test, * = *p* < 0.05, ** = *p* < 0.01, *** = *p* < 0.001, **** = *p* < 0.0001, ns = not significant. (D) Synergy maps and synergy scores showing multi-dose combination effects of DCZ0415 and JH-RE-06 on clonogenic survival of AsPc-1 and Capan-1 cells. (E) Immunoblots showing expression levels of the indicated DDR markers in control and TRIP13 siRNA-treated PDAC cell lines at different times after treatment with JH-RE-06 or Gemcitabine.

As expected from our studies with genetic ablation of *TRIP13*, the pharmacological TRIP13 inhibitor (DCZ0415) also synergized with JH-RE-06 or gemcitabine to kill HR-sufficient PDAC cell lines (Fig. 6D, Supplementary Figure S8). However, in the HR-deficient CAPAN-1 PDAC cell line, we observed no synergy or mild antagonism between DCZ0415 and JH-RE-06 or gemcitabine. Therefore, we conclude that the sensitivity of TRIP13-ablated (or TRIP13 inhibitor-treated) cells to JH-RE-06 and gemcitabine is due to reduced HR activity.

The lethality of therapeutic agents to HR-compromised cells has been attributed to both DNA DSB and ssDNA gaps (79). The extent to which each DNA lesion contributes to cell death remains heavily debated and may differ depending on biological context. To understand the molecular underpinnings of JH-RE-06 and gemcitabine-sensitivities in the setting of TRIP13-depleted PDAC we measured the accumulation of ssDNA and DSB markers in PDAC cells treated with JH-RE-06 or gemcitabine.

In HR-sufficient AsPC-1 and Pa03C cells, JH-RE-06 treatment induced accumulation of the DSB marker pATM (S1981) only after TRIP13-depletion (Fig. 6C). By contrast, in HR-deficient CAPAN-1 cells, pATM (S1981) was induced to high levels by JH-RE-06 treatment in both control and TRIP13-depleted cultures (Fig. 6C). These results suggest that HR-defects resulting from TRIP13-depletion (in AsPC-1 or Pa03C cells) or from intrinsic HR-defects (in CAPAN-1 cells) create a dependency on TLS for eliminating ssDNA gaps and averting DSB formation. Interestingly, in immunofluorescence microscopy analyses, we observed that TRIP13 knockdown led to reduced levels of nuclear 53BP1 foci, both in untreated and in JH-RE-06 treated cells (Supplementary Figure S7B). This result suggests that TRIP13 may also promote NHEJ in PDAC cells and that the synergistic lethality caused by TRIP13 loss in combination with JH-RE-06 may be driven by multiple mechanisms including HR and NHEJ defects.

The patterns of compensatory DSB signaling induced by gemcitabine in TRIP13-depleted cells were very different from those induced by JH-RE-06. Gemcitabine induced similar levels of pATM (S1981) in control and TRIP13-depleted cultures of AsPC-1 and Pa03C cells (Fig. 6C). However, in HR-deficient CAPAN-1 cells, gemcitabine induced a larger increase in pATM (S1981) levels in TRIP13-depleted cultures when compared with control cultures (Fig. 6C). We observed similar patterns for chromatin-binding of ZRANB3 and PRIMPOL (which mediates Template Switching and repriming in response to replication fork stalling) in CAPAN-1 cells: gemcitabine-induced levels of both ZRANB3 and PRIMPOL on chromatin increased more in TRIP13-depleted cells than in control (TRIP13-replete) cultures. These results indicate that the role of TRIP13 in promoting tolerance of gemcitabine-induced DNA lesions is not mediated solely via HR.

## DISCUSSION

Oncogene-induced DNA replication stress has been studied using a limited number of representative oncogenes such as *CCNE1*, typically expressed at very high levels in pathologically-irrelevant cell lines such as U2OS or fibroblasts. Here we have established a new preneoplasia model in which oncogenic KRAS is expressed at pathologically-relevant levels in precursor cell line that is appropriate for PDAC. Using this more pathologically-relevant model system, we find that low level KRAS expression is mitogenic, yet does not significantly induce replication stress, senescence, or major disruptions (such as the remarkable whole genome duplication which results when *CCNE1* is massively overexpressed in U2OS cells (29). Using a comparison of our dox-inducible KRAS^G12V^ model and KRAS-mutant PDAC patient RNA-seq data followed by a candidate gene approach, we identified TRIP13 as a new mediator that allows HPNE to tolerate the DNA replication stress caused by low-level oncogenic KRAS.

TRIP13 targets multiple protein complexes in the DDR and cell cycle including REV7-Shieldin, REV7-TLS, REV7/MAD2B-APC, and likely others. In the setting of untransformed HPNE cells expressing oncogenic KRAS, the role of TRIP13 in sustaining DNA synthesis and viability is most readily explained by its ability to promote HR. This conclusion is suggested both by the epistatic relationship between TRIP13 and HR genes for tolerating *KRAS^G12V^* expression, and by our observation that TRIP13-ablation in KRAS^G12V^-expressing HPNE induces a hallmark HRD phenotype, namely PARPi-sensitivity. In addition to PARPi-sensitivity, HRD can create a dependency on TLS for gap-filling (80) – a feature that fully explains JH-RE-06-sensitivity of TRIP13-depleted cells that express KRAS^G12V^. HR contributes to tolerance of replication stress induced by other oncogenes including CCNE1 (81) and β-catenin (27). Moreover, TRIP13 is overexpressed in many cancers (82). Therefore, it will be interesting to determine whether TRIP13 induction in malignant cells is a general adaptive response to diverse forms of oncogene-induced DNA replication stress.

In addition to its role in allowing neoplastic cells to tolerate KRAS^G12V^ (and perhaps other oncogenes), we show that TRIP13 also allows PDAC cells to tolerate therapeutic agents including the PARP inhibitor Olaparib.

Interestingly however, the impact of TRIP13 on PARPi-sensitivity appears to be context-dependent. For example, we show that TRIP13-knockdown does not increase olaparib-sensitivity of HR-deficient Capan-1 PDAC cells, indicating that in these cells the role of TRIP13 in resisting PARP inhibitors is HR-dependent. In contrast, TRIP13-depletion led to increased Olaparib-sensitivity in BRCA1-deficient SUM149PT breast cancer cells (83). Paradoxically, TRIP13 depletion did not affect Olaparib-sensitivity in wild-type RPE, yet TRIP13-loss (using shRNA- and sgRNA) rendered BRCA1-deficient RPE less sensitive to Olaparib (84). Most likely the variability of Olaparib-sensitivity phenotypes caused by TRIP13 loss reflects the pleiotropic roles of TRIP13 in DNA repair. Since DDR features can vary tremendously between different cancer cells, the mechanisms by which TRIP13 modulates the DDR are likely to vary significantly depending on cellular background.

In addition to the well-studied complexes of REV7 with REV3 (which promotes TLS) and shieldin (which is inhibitory for resection and HR), REV7 also associates with chromosome alignment-maintaining phosphoprotein 1 or CHAMP1 (85) to promote HR (86). If the relative pools of REV7 in complex with its various partners vary between different cell types, this could well explain how TRIP13-mediated dissociation of REV7 protein complexes has pleiotropic effects on HR in different cells.

It is also possible that the effects of TRIP13 on DDR are mediated by REV7-independent mechanisms. Some studies have found that TRIP13 promotes NHEJ directly (75,76). For example, TRIP13 interacts with DNA-PKcs complex proteins to stimulate NHEJ, even in HR-sufficient cells (75). TRIP13 also facilitates the the interaction between MDC1 and the MRN complex to promote and amplify ATM signaling (76). The REV7- related germ cell protein HORMAD1 is also a probable TRIP13 target, since it is removed from the synaptonemal complex of unsynapsed meiotic chromosomes in a TRIP13-dependent manner (87). HORMAD1 is a Cancer/Testes Antigen (CTA) that is mis-expressed in many cancer cells where it promotes HR (88,89).

Although the putative binding partner of HORMAD1 in promoting HR in cancer cells is unknown, it is likely that HR-stimulatory HORMAD1 protein complexes would also be dissociated by TRIP13. Since HORMAD1 expression in cancer cells is highly variable, TRIP13 could have very different effects on HR in different cancer cells depending on their HORMAD1 status. Similarly, variability in the availability or deployment of any TRIP13- interacting DDR factors and pathways might determine how TRIP13 inhibition affects DNA repair in different cancer cells While this work was in progress, two other studies demonstrated enrichment of TRIP13 expression in PDAC and other GI tumors when compared with normal tissues (82,90). Consistent with our study, Afaq and colleagues also showed that TRIP13-depletion led to reduced growth of PDAC cell lines cultured *in vitro* and when xenografted into mice (90). Moreover, Afaq et al. demonstrated that genetic or pharmacological ablation of TRIP13 led to increased gemcitabine-sensitivity in cultured cells and *in vivo* (90). Mechanistically, those workers suggested that TRIP13 sustains tumorigenic characteristics in PDAC by maintaining expression of FGFR4, phosphorylation of STAT1, and WNT/β-catenin signaling (90). It will be interesting to determine whether TRIP13 sustains tumorigenic signaling pathways (such as FGFR4, STAT1 and WNT/β-catenin) secondarily to its role in tolerating DNA replication stress, or instead whether protein complexes targeted by TRIP13 modulate mitogenic signaling directly independently of the DDR. Owing to its upregulated expression in cancer cells and its roles in sustaining neoplastic cells, and pharmacological tractability, TRIP13 remains a very attractive therapeutic target (91).

Since TRIP13 has pleiotropic roles in the DDR and other processes, it is necessary to determine how the various TRIP13 effector pathways function in cancer cells. Elucidating how TRIP13-interacting pathways are wired in different cancer settings may reveal opportunities for precision medicine - for example by identifying those tumors that are most sensitive to TRIP13 inhibitors alone or in combination with other agents (such as PARP or TLS inhibitors).

## DATA AVAILABILITY

RNA-seq data for hTERT-HPNE cells underlying this article are available in BioProject ID PRJNA1137433. The TCGA pancreatic adenocarcinoma (TCGA-PAAD) data was retrieved using TCGAbiolinks R package (version 2.30.0) and is available on TCGA portal (https://portal.gdc.cancer.gov/). Unnormalized counts and differential gene expression statistics are available in Supplementary Table 2. Steps, criteria and code used for HPNE differential gene expression analysis, TCGA-PAAD analysis, and PHATE analysis using 4i experiment are available in the GitHub repository: https://github.com/vazirilb/PancreaticCancer-DifferentialExpression-4iPHATE.

## SUPPLEMENTARY DATA

Supplementary Data are available at NAR online

## ACKNOWLEDGEMENTS

We thank Dr. Pablo Ariel for his assistance in microscopy image acquisition. Microscopy was performed at the Microscopy Services Laboratory at the University of North Carolina, Chapel Hill, which is supported in part by P30 CA016086 Cancer Center Core Support Grant to the UNC Lineberger Comprehensive Cancer Center.

The Andor Dragonfly microscope was funded with support from National Institutes of Health grant S10OD030223. Research reported in this publication was supported in part by the North Carolina Biotech Center Institutional Support Grant 2012-IDG-1006. Animal studies were performed within the UNC Lineberger Preclinical Research Unit at the University of North Carolina at Chapel Hill which is supported in part by an NCI Center Core Support Grant (CA16086) to the UNC Lineberger Comprehensive Cancer Center.

