## Supplementary Figures for "TRIP13 protects pancreatic cancer cells against intrinsic and therapy-induced DNA replication stress"

**List of Supplementary Figures**

**Content**

**Figures S1-S9 Page 2-17**

**Figure S1.**

(A) Flow cytometry profiles showing PI staining and levels of incorporated BrdU in HPNE empty vector cells 2 days after infection Cyclin E1 adenovirus (or with an 'empty vector' control) adenovirus. (B)-(F) Images showing microscopy analysis of H&E-stained sections of pancreata from experimental mice orthotopically injected with the following cell lines (and conditionally given Dox in their drinking water as indicated): Pa02c, no Dox (B); HPNE empty vector (EV), no Dox (C); HPNE EV, +Dox (D); HPNE KRAS$^{G12V}$, no Dox (E); HPNE KRAS$^{G12V}$, +Dox (F). Scale: 2x =500 mm; 10x = 100 mm. (G) Flow cytometry gating strategy

**Figure S2.**

(A) Z-normalized PHATE plots of tested proteins colored by the signal intensity of each cell cycle and DNA damage markers in empty vector (EV), empty vector + 200 ng/mL Dox (EV Dox), KRAS$^{G12V}$and KRAS$^{G12V}$ + 200 ng/mL Dox (KRAS$^{G12V}$ Dox) after 3 days of mutant KRAS induction.

**Figure S3.**

Z-normalized protein (4i) signals of yH2AX (S139), phosphorylated-TP53, p21, phosphorylated-p21 (T145) and phosphorylated-p27 (T157) in G1, S and G2/M cell cycle phases of empty vector and KRAS$^{G12V}$-expressing HPNE and their relevant controls after 3 days of Dox-treatment.

**Figure S4.** siRNA screen of select upregulated DDR genes. Graphs summarizing (A) colony survival, (B) distribution of cells between different cell cycle phases, and (C) percentage BrdU incorporation for siRNA knockdown of DDR genes in the screen including controls KRAS and TP53.

**Figure S5.** (A) Immunoblots showing Cyclin E and MYC overexpression levels in HPNE cells. (B) Immunoblots showing TRIP13 knockdown. (C) Graph showing effect of TRIP13 knockdown on numbers of viable HPNE cells after 2 days of Cyclin E1 or Myc overexpression. (D) Graph showing quantification of BrdU-incorporation after 2 days of Cyclin E1 or Myc overexpression. Error bars represent standard deviation (SD) for biological duplicates. Statistical analysis: two-way ANOVA followed by Tukey's multiple comparisons test, * = $p < 0.05$, ** = $p < 0.01$, *** = $p < 0.001$, **** = $p < 0.0001$, ns = not significant. (E) Plot comparing TRIP13 expression between different genders in TCGA-PAAD.

**Figure S6.** Analysis of DNA replication dynamics showing effects of TRIP13 knockdown on levels of stalled DNA replication forks (A) and new replication origin firing (B) in HPNE cells. Error bars represent standard deviation (SD) for biological triplicates. Statistical analysis: two-way ANOVA followed by Tukey's multiple comparisons test, * = $p < 0.05$, ** = $p < 0.01$, *** = $p < 0.001$, **** = $p < 0.0001$, ns = not significant.

**Figure S7.** (A) Images of representative HPNE nuclei showing patterns of 53BP1 and DAPI staining for the indicated experimental conditions (Left Panel) and quantification of 53BP1 foci-containing cells for each experimental condition (Right Panel). Error bars represent standard deviation (SD) for biological duplicates. Statistical analysis: two-way ANOVA followed by Tukey's multiple comparisons test, * = $p < 0.05$, ** = $p < 0.01$, *** = $p < 0.001$, **** = $p < 0.0001$, ns = not significant. (B) Images of representative AsPc1 nuclei showing patterns of 53BP1 and DAPI staining under the indicated experimental conditions (Left Panel) and Quantification of 53BP1 foci-containing cells under each experimental condition (Right Panel). Error bars represent standard deviation (SD) for biological duplicates. Statistical analysis: two-way ANOVA followed by Tukey's multiple comparisons test, * = $p < 0.05$, ** = $p < 0.01$, *** = $p < 0.001$, **** = $p < 0.0001$, ns = not significant.

**Figure S8.** TRIP13 inhibition is synergistic with therapeutic agents. (A) Synergy maps and synergy scores showing multi-dose combination effects of DCZ0415 and JH-RE-06 on clonogenic survival of AsPc-1, Panc10.05, Pa02c, Pa03c and Capan-1 cells.

**Figure S9.** Immunoblots showing the effect of $KRAS^{G12V}$-induction on ERK phosphorylation and RAD51 expression.

Supplementary Figure S1

A

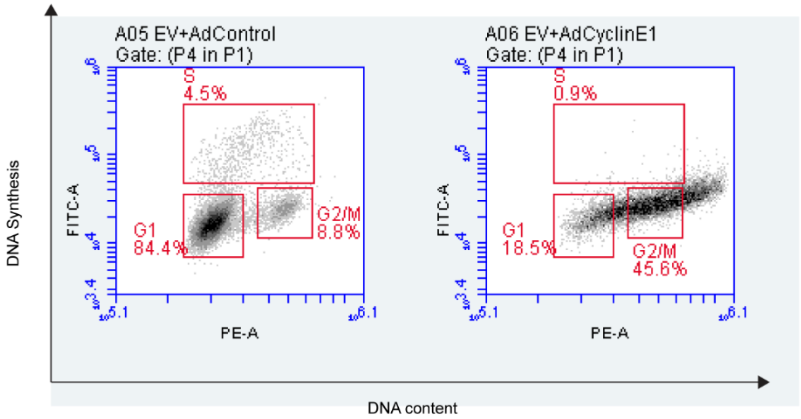

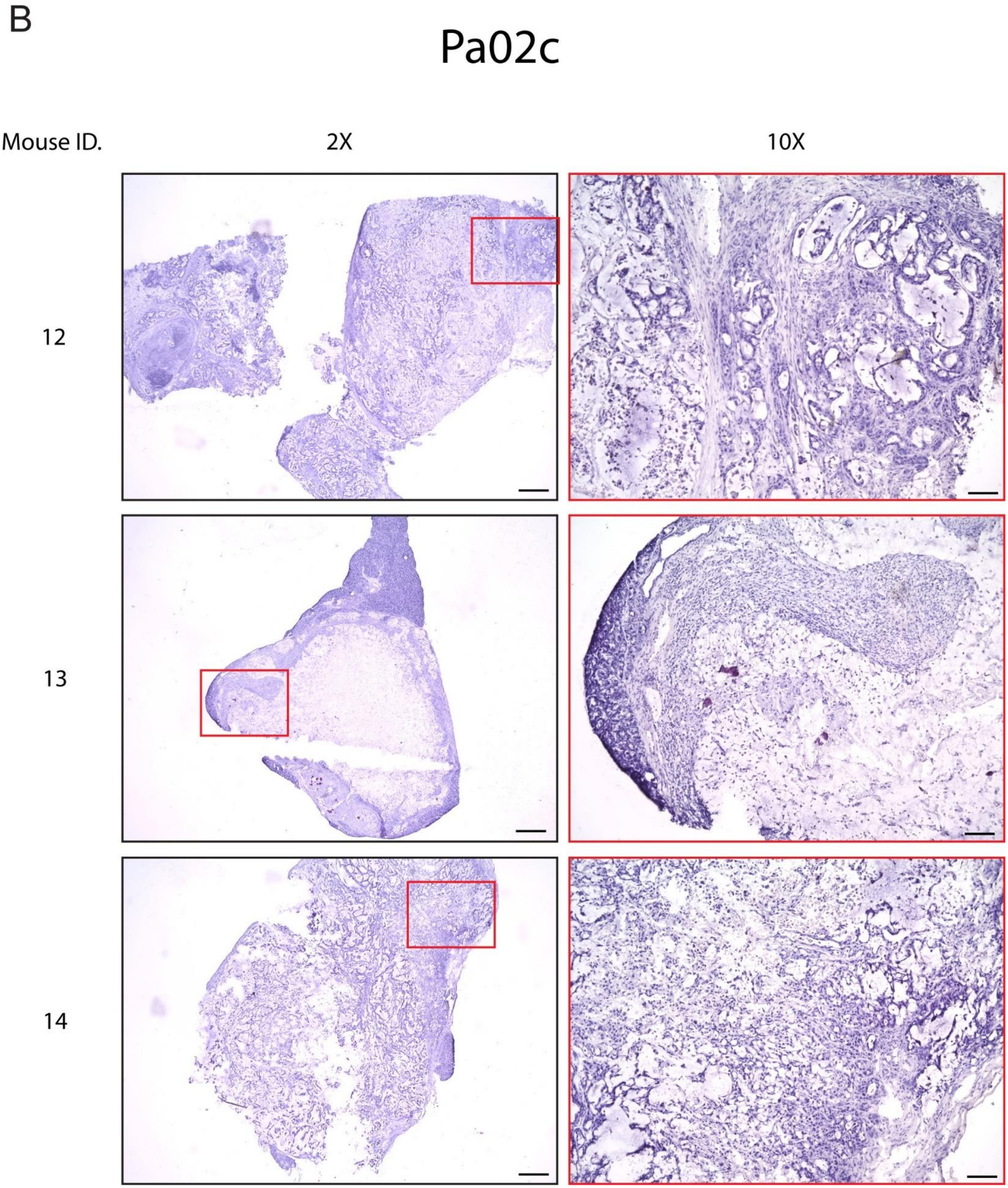
B
Pa02c
Mouse ID.
2X
10X
12
13
14

Supplementary Figure S1

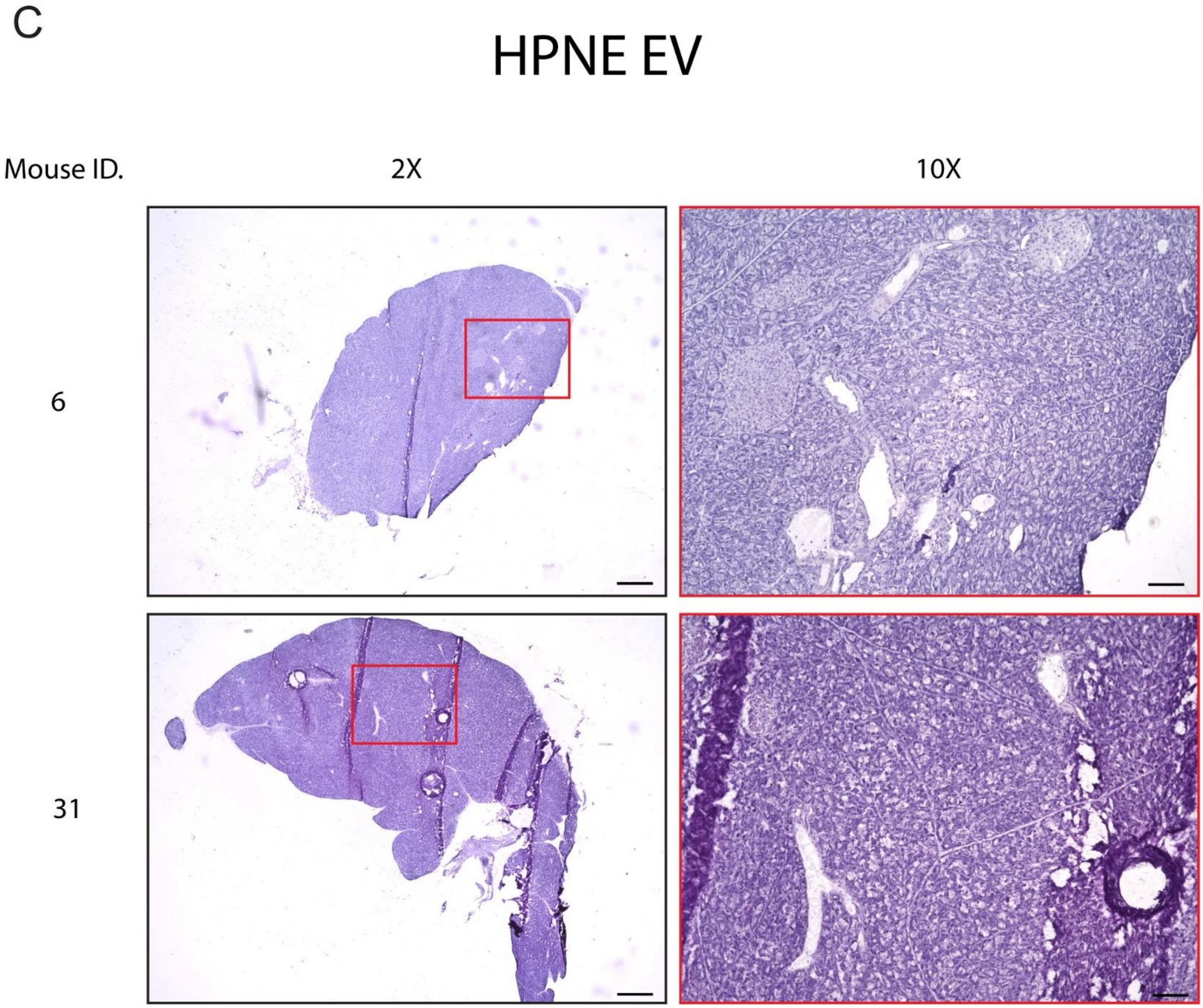

Supplementary Figure S1

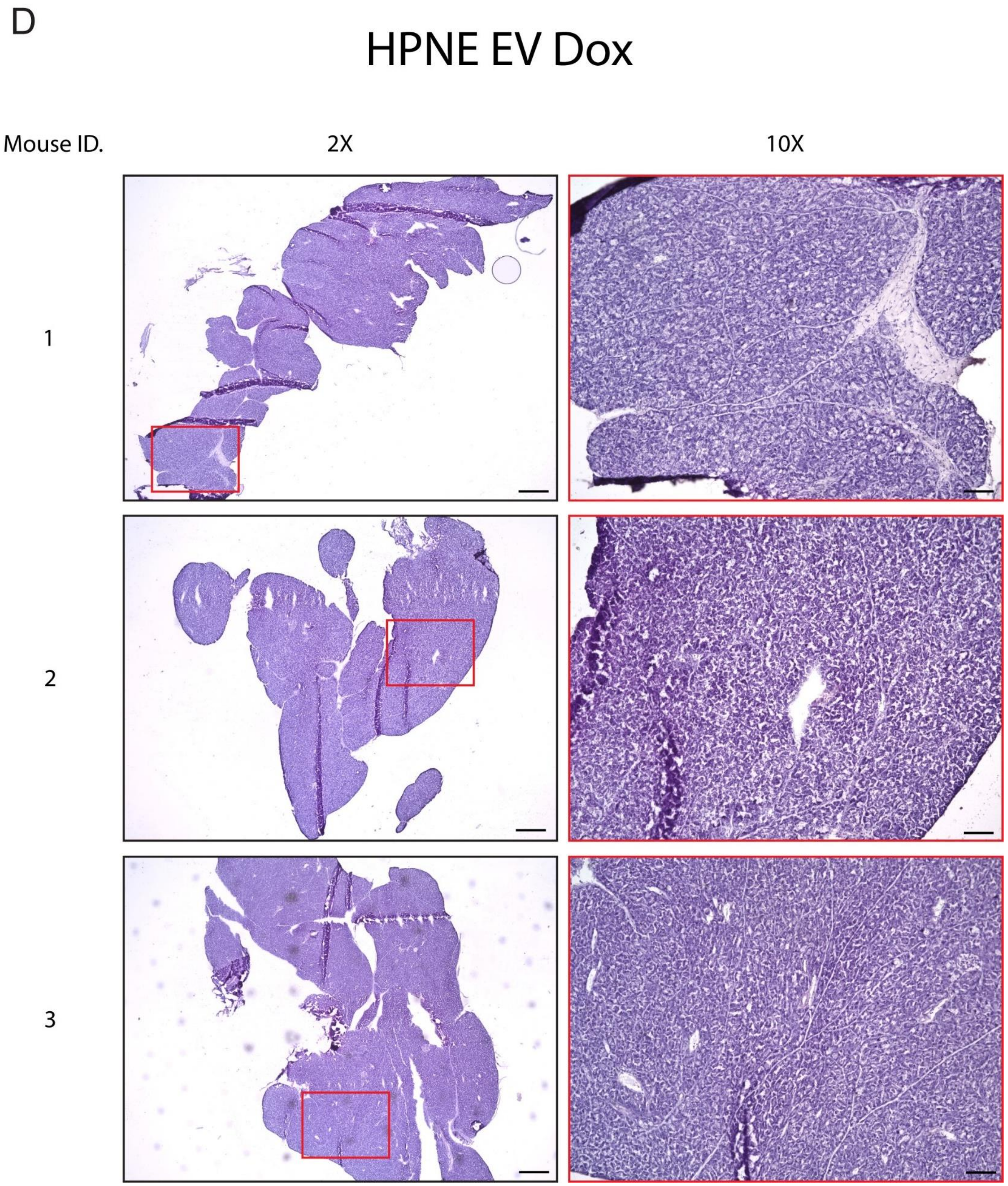

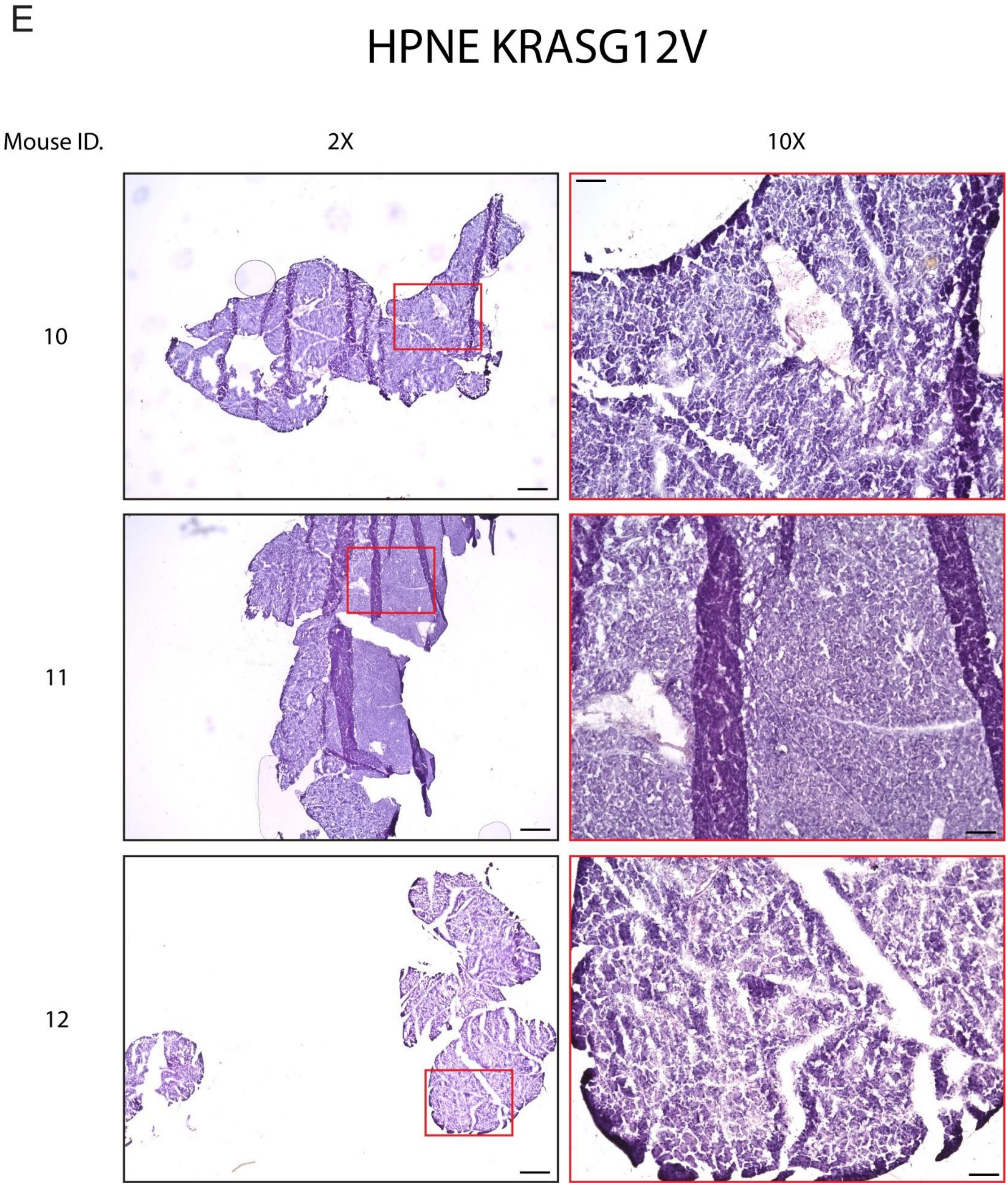
E
HPNE KRASG12V
Mouse ID.
2X
10X
10
11
12

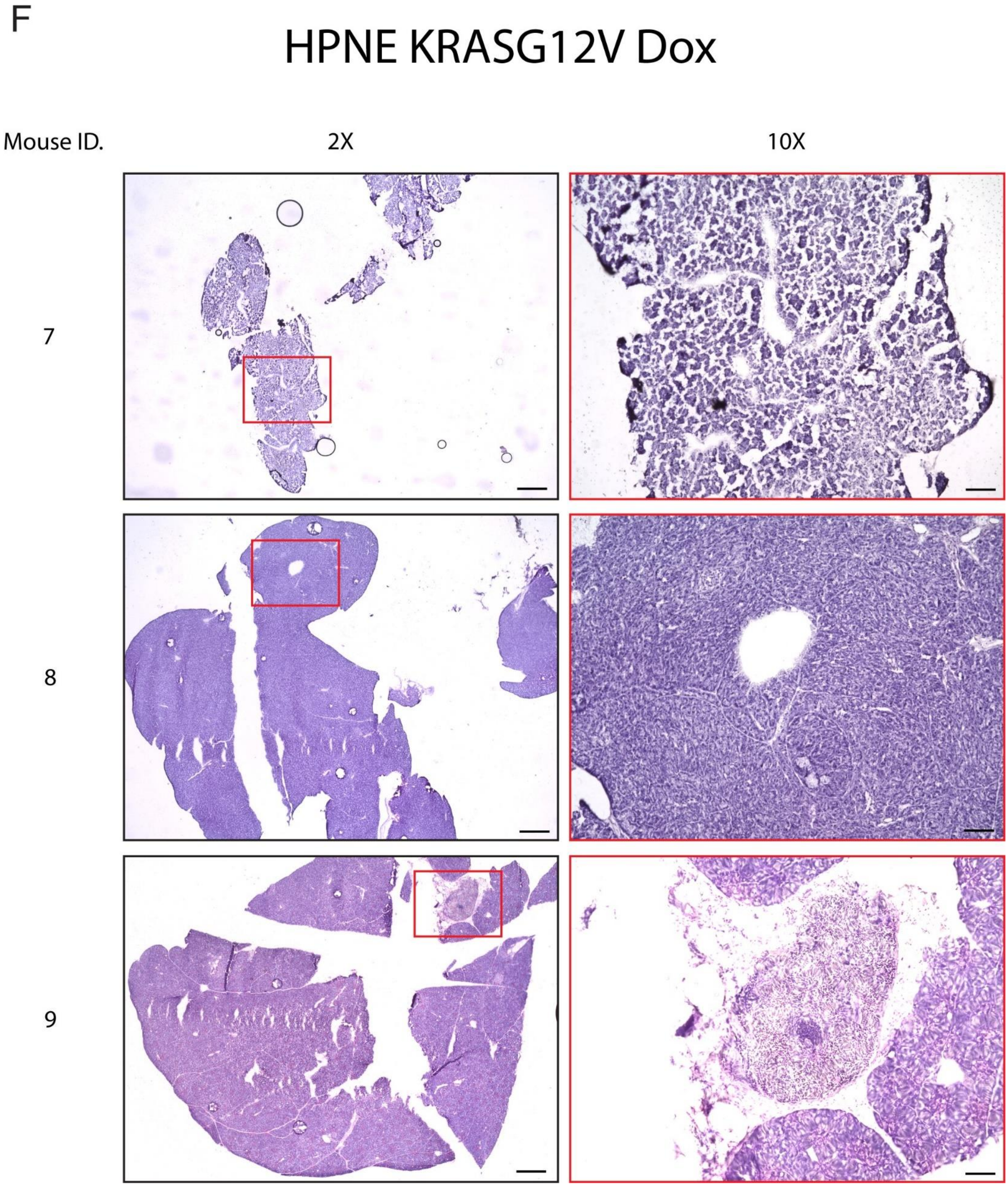
F
HPNE KRASG12V Dox
Mouse ID.
2X
10X
7
8
9

Supplementary Figure S1

G

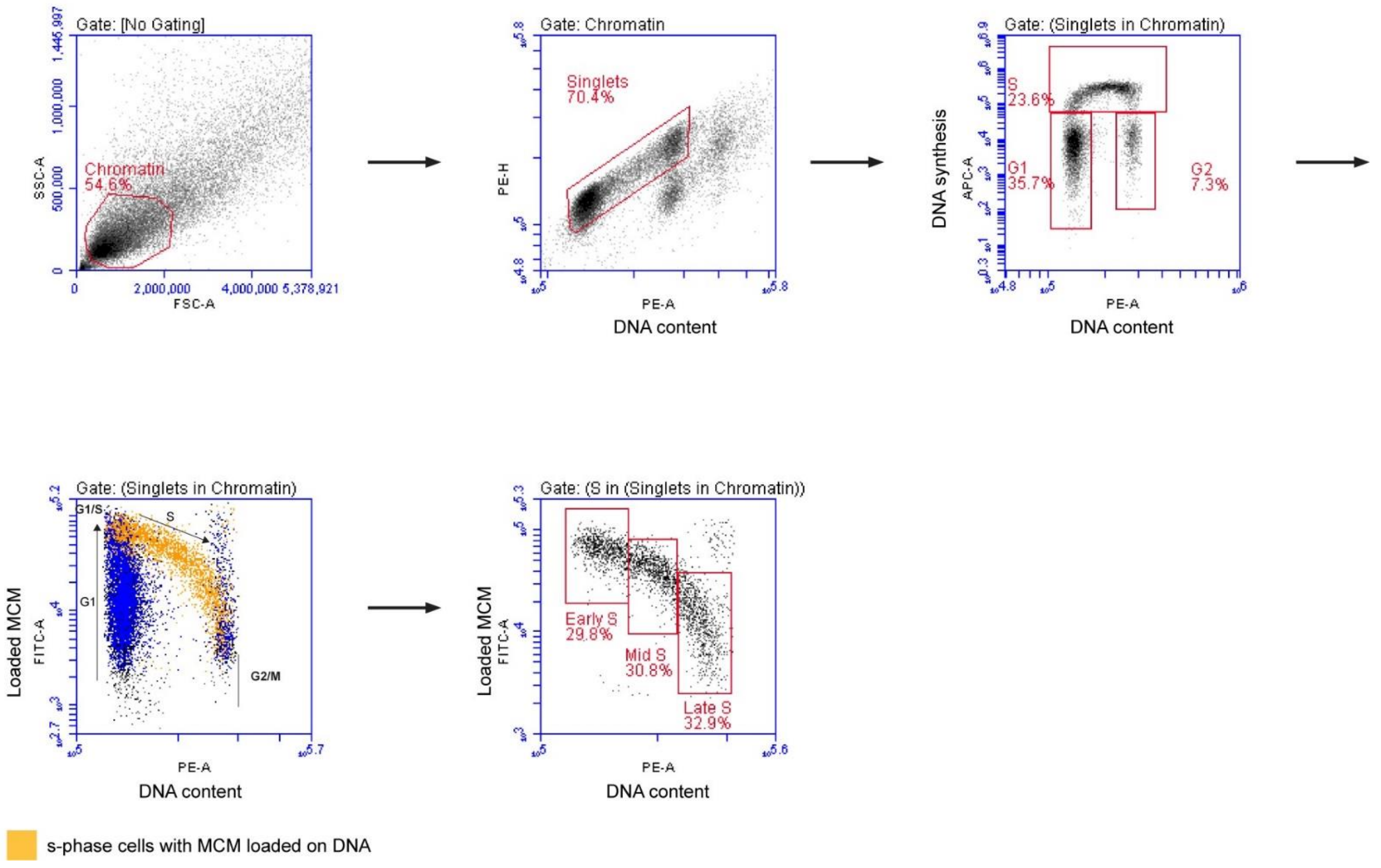

Supplementary Figure S2

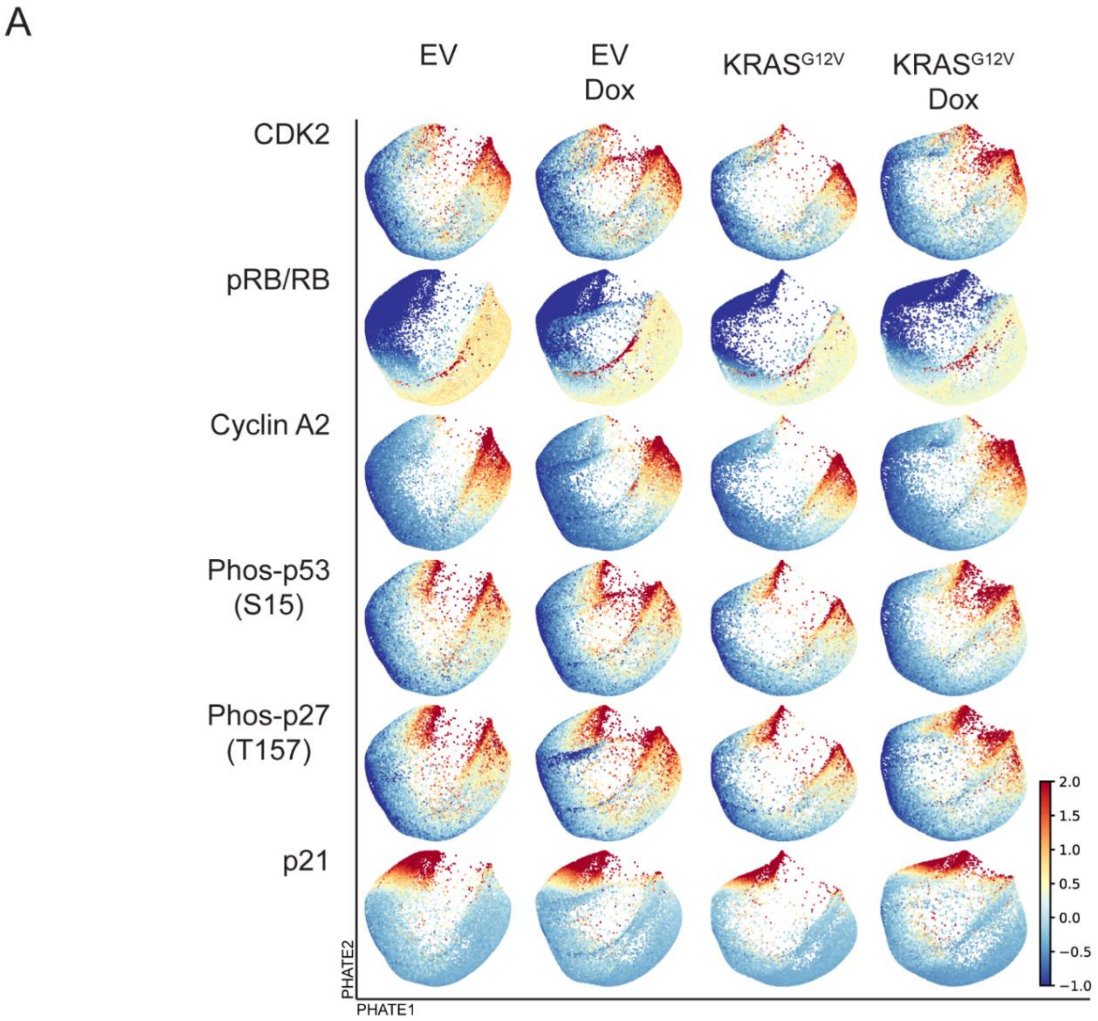

Supplementary Figure S3

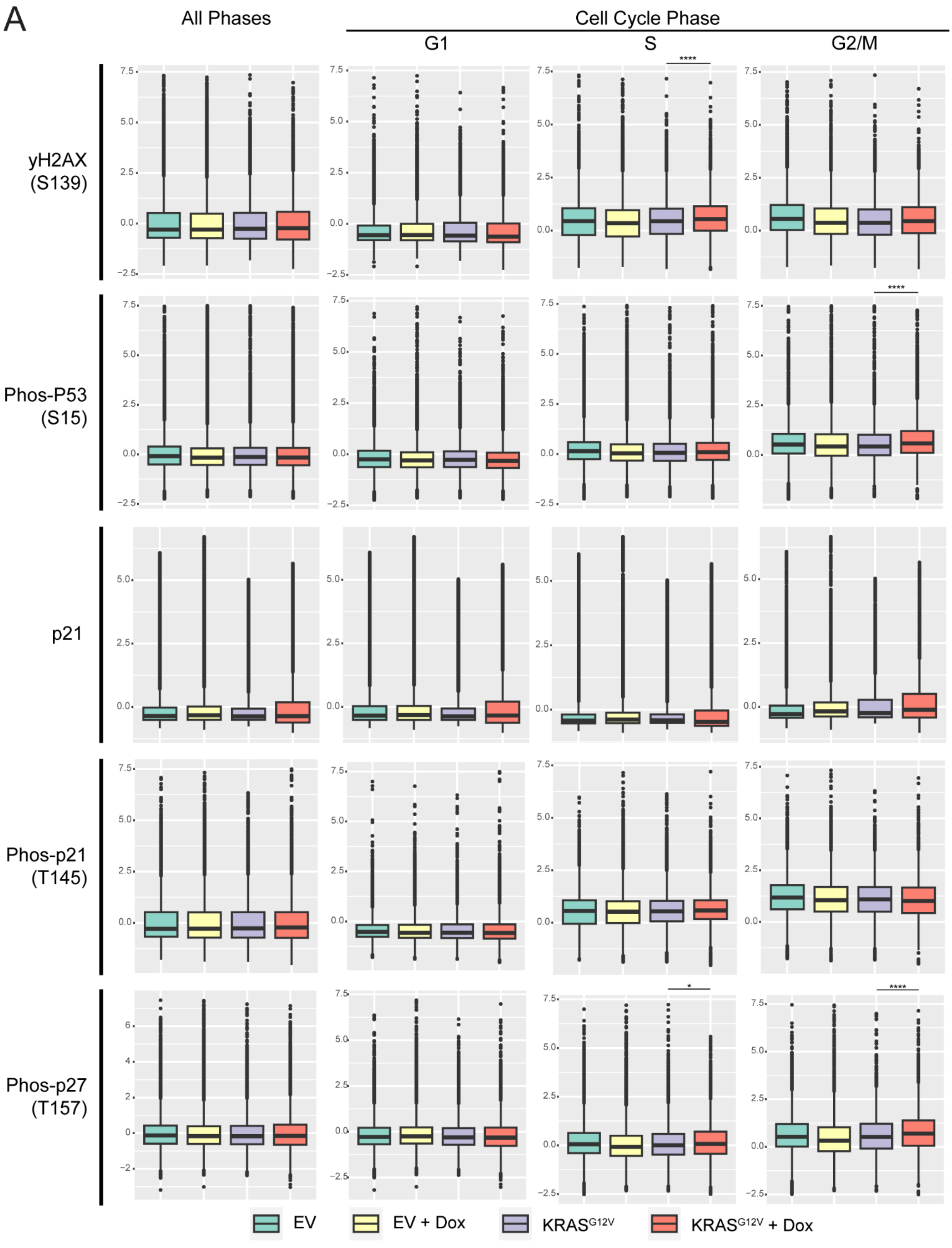

Supplementary Figure S4

A

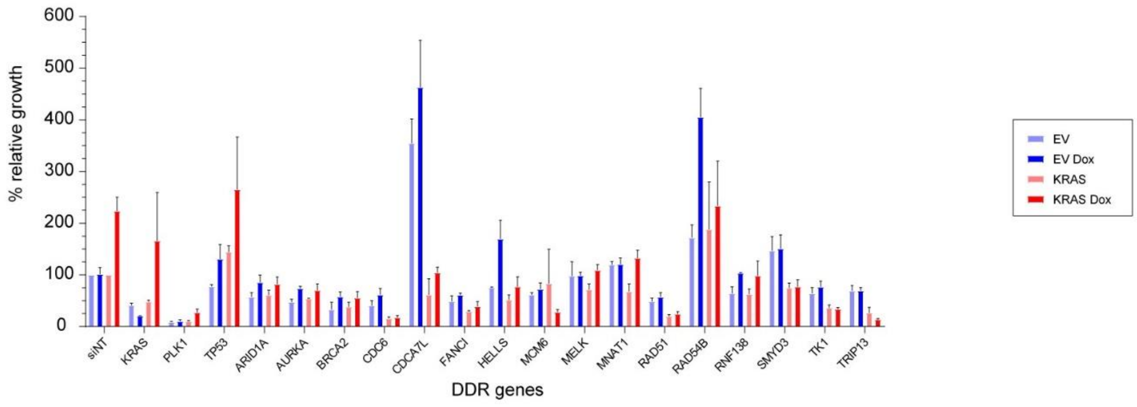


B

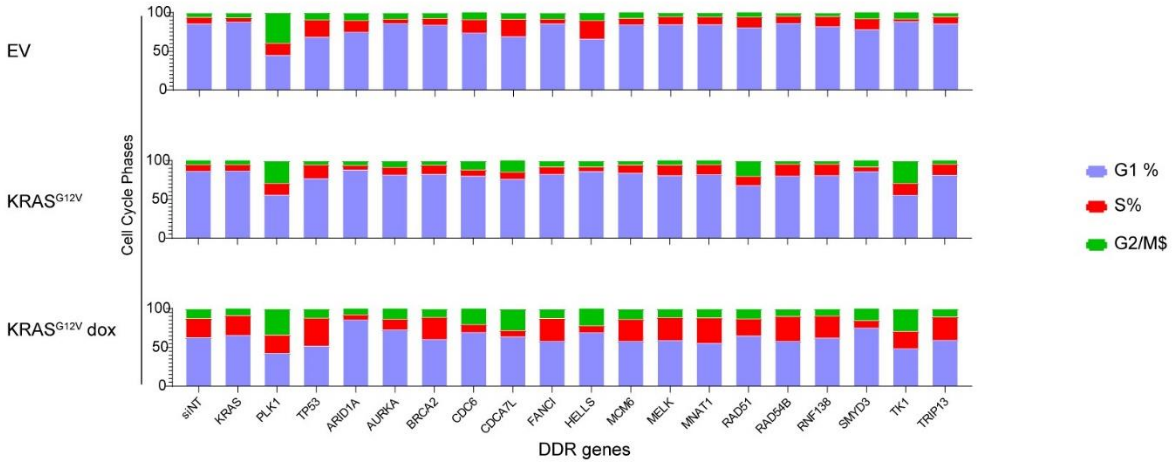


C

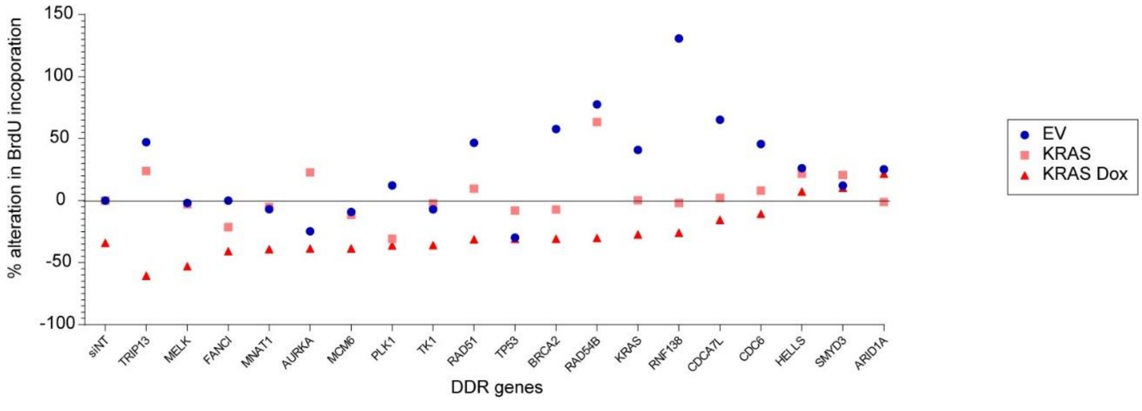

Supplementary Figure S5

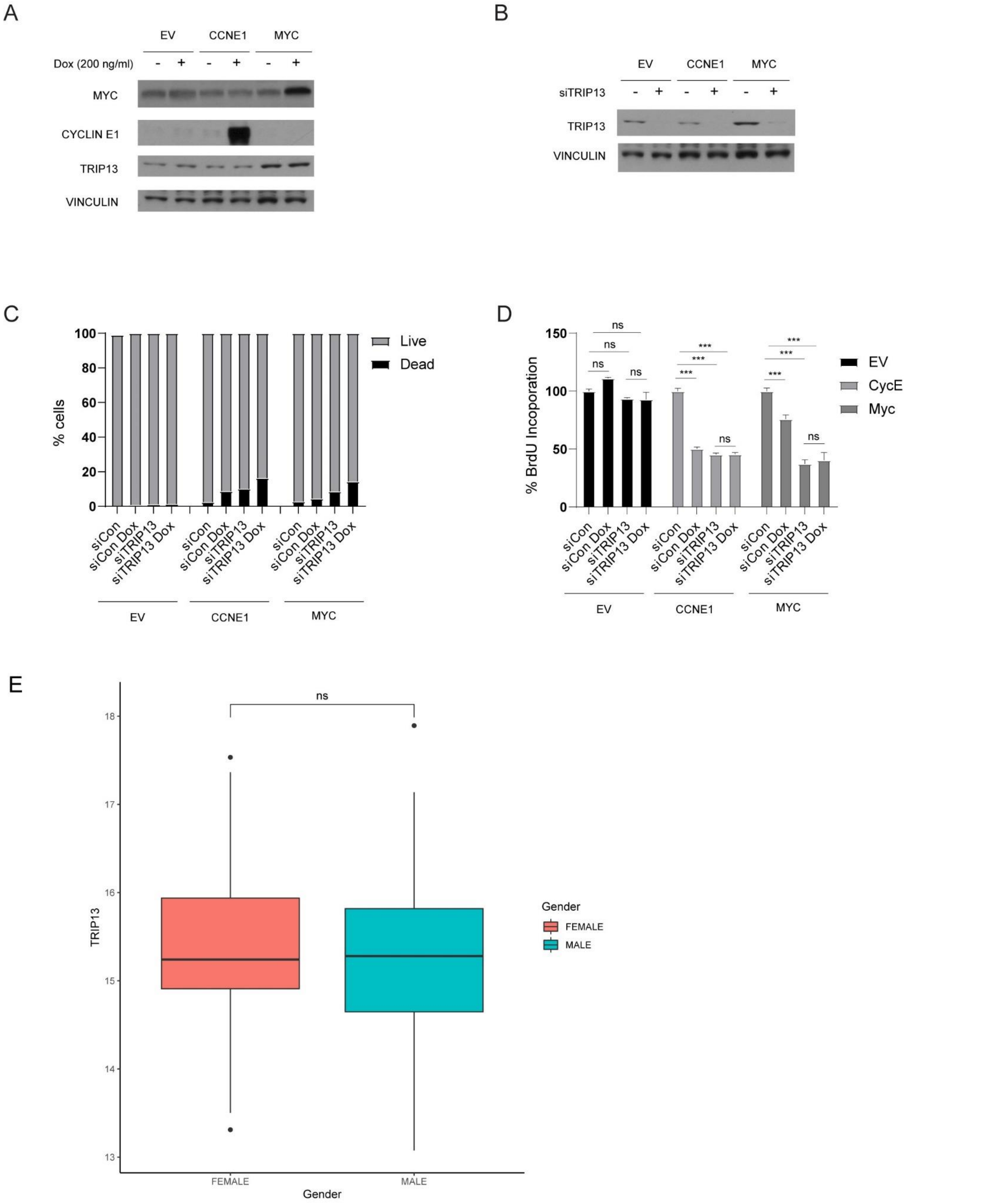

Supplementary Figure S6

A

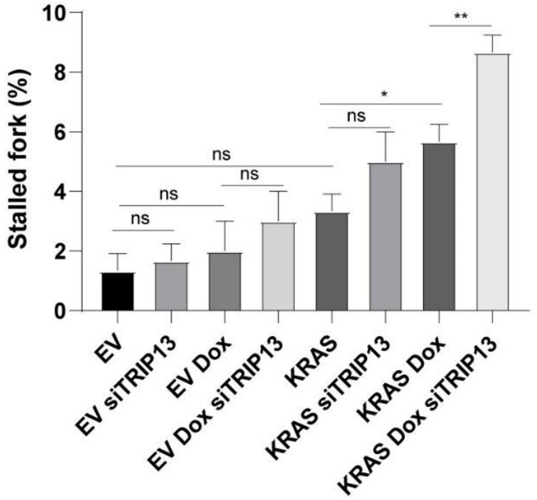


B

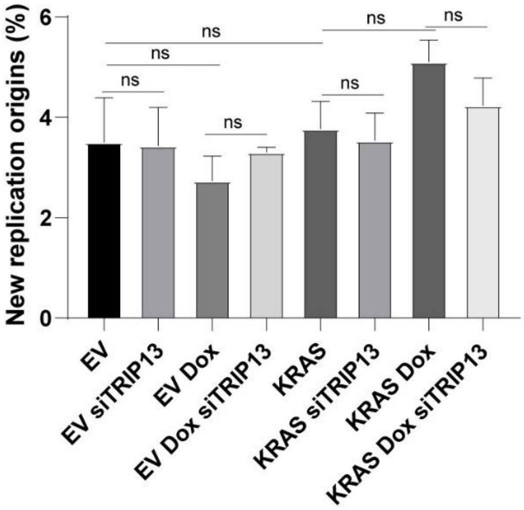

Supplementary Figure S7

A

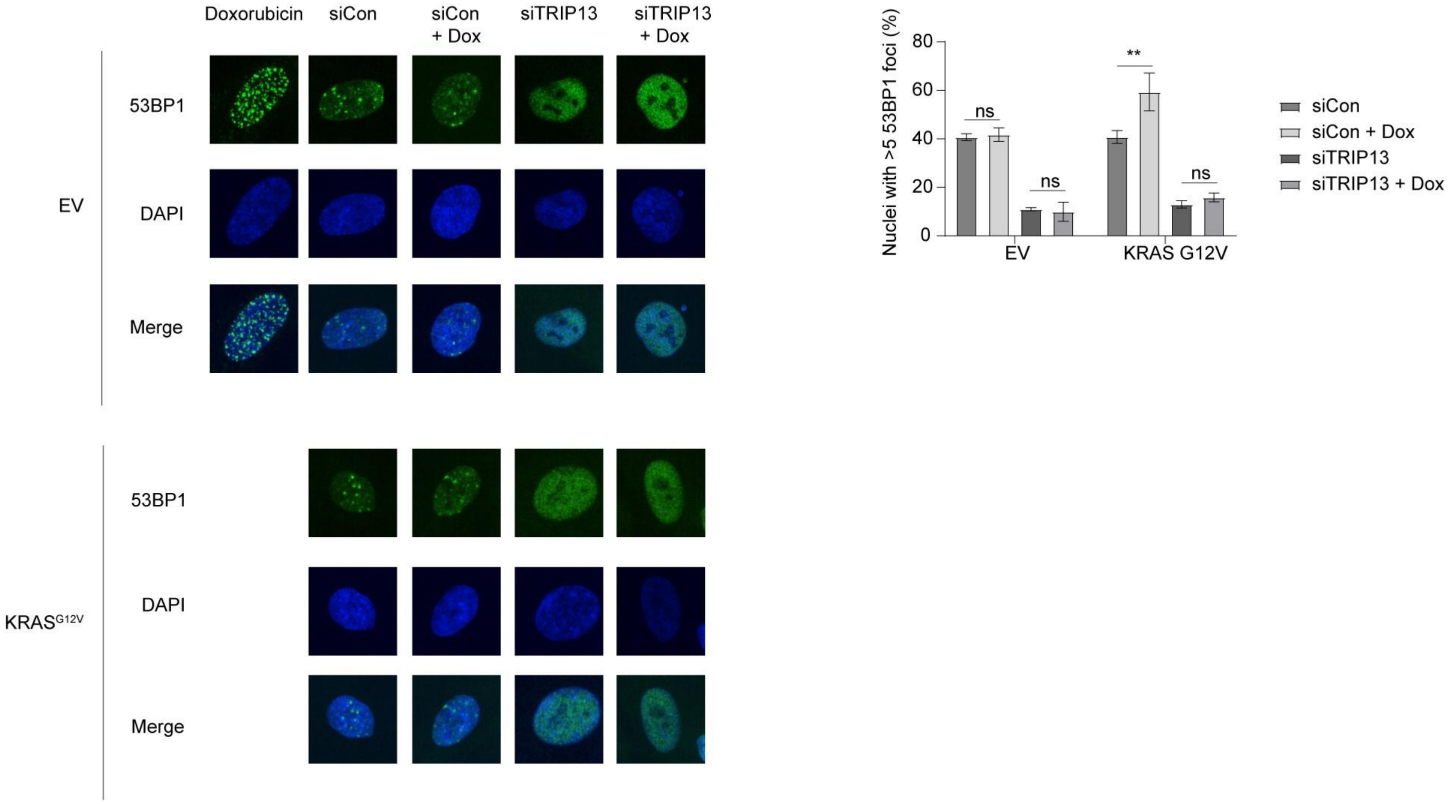


B

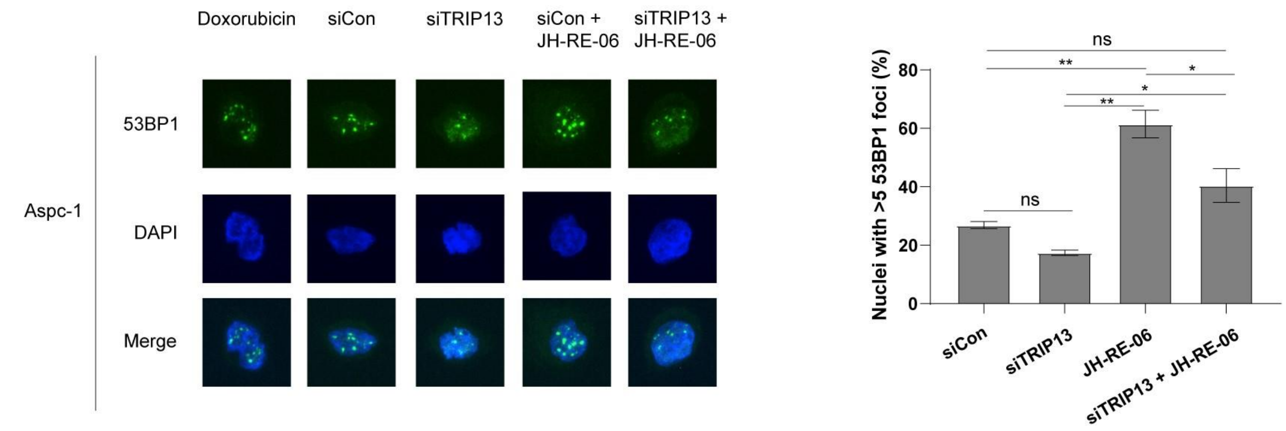

Supplementary Figure S8

A

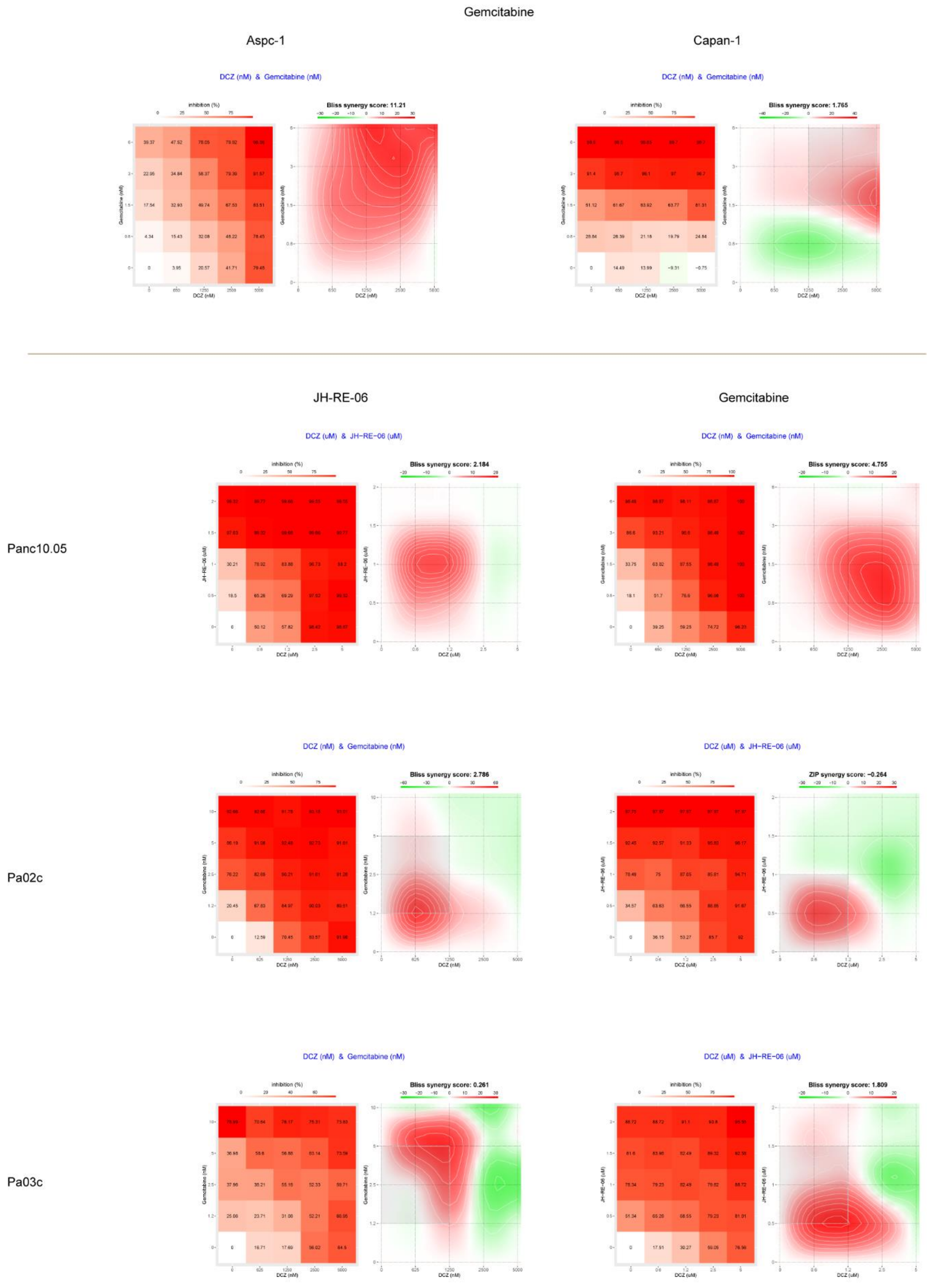

Supplementary Figure S9

A

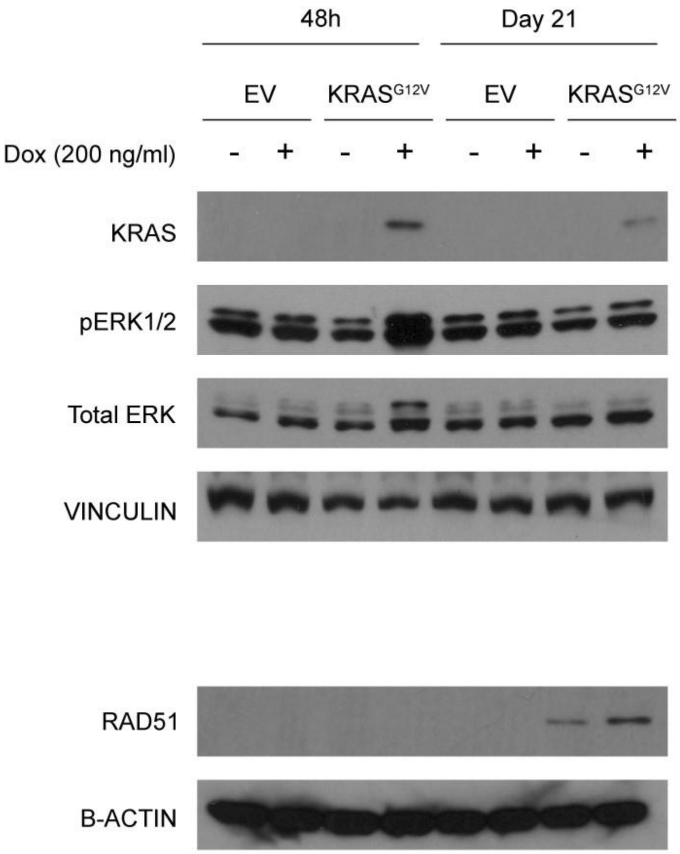
